# Neurons undergo pathogenic metabolic reprograming in models of familial ALS

**DOI:** 10.1101/2021.08.20.457111

**Authors:** Sean-Patrick Riechers, Jelena Mojsilovic-Petrovic, Mehraveh Garjani, Valentina Medvedeva, Casey Dalton, Gerald Dienel, Robert G. Kalb

## Abstract

Normal cellular function requires a rate of ATP production sufficient to meet demand. In most neurodegenerative diseases (including Amyotrophic Lateral Sclerosis, ALS), mitochondrial dysfunction is postulated raising the possibility of impaired ATP production and a need for compensatory maneuvers to sustain the ATP production/demand balance. We find in our rodent models of familial ALS (fALS), impairment in neuronal glycolytic flux with maintained or enhanced activity of the citric acid cycle. This rewiring of metabolism is associated with normal ATP levels and redox status, supporting the notion that mitochondrial function is not compromised in neurons expressing fALS genes. Genetic loss-of-function manipulation of individual steps in the glycolysis and the pentose phosphate pathway blunt the negative phenotypes seen in various fALS models. We propose that neurons adjust fuel utilization in the setting of neurodegenerative disease-associated mitochondrial dysfunction in a baleful manner and targeting this process can be healthful.

## Introduction

Alterations in mitochondrial structure and function are commonly observed in adult-onset neurodegenerative diseases (Archer, 2013; Area-Gomez et al., 2019; Casajus Pelegay et al., 2019; Cowan et al., 2019; Gao et al., 2019; Lin and Beal, 2006; Rangaraju et al., 2019; Wu et al., 2019). In ALS, mitochondrial dysfunction impairs the efficiency of electron transport chain (ETC) activity and ATP production (Mattiazzi et al., 2002) (Bowling et al., 1993; Dupuis et al., 2004; Jung et al., 2002; Magrané et al., 2012), and leads to accumulation of reactive oxygen and nitrogen species (Greco et al., 2019; Kim et al., 2015; Pirie et al., 2021), abnormal handling of intracellular calcium, and cytochrome C release and apoptosis (Smith et al., 2019). The extent to which these alterations in mitochondrial function impair cellular operations is unclear.

Therapeutic intervention based on combating these mitochondrial abnormalities have displayed variable success in mouse models of ALS and humans (Wu et al., 2019) (Marques and Wyse, 2019), reviewed in Vandoorne et al. (Vandoorne et al., 2018).

Eukaryotic cells monitor their energy economy very carefully to ensure energy production and energy utilization are matched (Dienel, 2019; Hardie et al., 2012; Mihaylova and Shaw, 2011). From the energy budget perspective, synaptic transmission, action potentials and the maintenance of the resting membrane potential are considered the most energetically expensive neuronal processes (Yu et al., 2018). Even a modest, chronic decrement in ATP production could adversely affect fundamental neuronal biology and consequently impair neuronal functions and survival. In addition, on a vastly different time scale, the activity-driven local ATP synthesis is required for moment-to-moment synaptic function (Licznerski et al., 2020; Mann et al., 2021; Rangaraju et al., 2014). Neuronal glucose oxidative phosphorylation is the prime source of ATP subserving pre- and post-synaptic element cell biology (Hall et al., 2012). If neuronal mitochondrial dysfunction in the setting of ALS significantly upset the balance of ATP production with ATP utilization it would negatively impact all cellular functions.

To achieve cellular homeostasis in the setting of neurodegenerative disease-associated mitochondrial dysfunction, neurons could reduce anabolic processes and/or attempt to re-wire metabolism to maintain adequate ATP levels. Two general strategies could be employed to increase ATP production. First, maneuvers could be deployed to increase NADH production by the tricarboxylic acid (TCA) cycle, forcing more electrons into the ETC. Second, glycolysis, might be stimulated. While a less efficient method for ATP production, work in the cancer field establishes that aerobic glycolysis can be a substantial energy source (DeBerardinis and Chandel, 2016). In either scenario, re- wiring metabolism could have unintended consequences on cellular biochemistry. For example, glycolytic intermediates are the starting point for the pentose phosphate pathway (PPP, which generates NADPH, essential for glutathione and thioredoxin control of reactive oxygen species) (Kletzien et al., 1994; Stanton, 2012; Wood, 1986), the hexosamine biosynthetic pathway (HBS, which generates UDP-N-acetylglucosamine, essential for N- and O-glycans within the endoplasmic reticulum) (Denzel et al., 2014; Slawson and Hart, 2011; Wang et al., 2014) and the synthesis of amino acids. Enhanced glycolytic flux, at the expense of PPP or HBS, might adversely impact redox status, ER stress and protein synthesis. In addition, the source of ATP (that is, glycolysis versus ETC) has cell type specific effects on signaling pathways. For example, immune cell Akt-FOXO1 signaling is driven by mitochondrial derived ATP in naïve T cells and by glycolysis in effector T cells (Xu et al., 2021).

The cardinal feature of ALS is degeneration of upper and lower motor neurons and thus understanding the contribution of metabolic dysfunction of neurons is a critical issue.

To understand how fuel is utilized we undertook glucose isotopologue investigations in cultured rodent cortical neurons expressing two different fALS genes – Cu^++^/Zn^++^ superoxide dismutase (SOD1) or TAR DNA binding protein of 43 kDa molecular weight (TDP43). Based on our interpretation of these metabolite isotopologue results, we hypothesized that altering the expression of individual glycolytic genes could modify phenotypes evoked by expression of wild type (“WT”) or disease-causing missense versions (“mutant” or “m”) of SOD1 or TDP43. We tested this hypothesis in a series of functional studies employing *Saccharomyces cerevisiae, Caenorhabditis elegans* and *Rattus norvegicus* spinal cord neuron platforms. These integrated investigations reveal that: 1) There is little evidence of neuronal mitochondrial dysfunction in our models when focusing on outputs such as energy charge or redox state and 2) manipulating fuel utilization by neurons can blunt neuronal dysfunction and death in a variety of ALS models.

## Results

### Metabolomic investigations of cortical neurons expressing ALS-causing mutant proteins

For metabolomic studies, neuronal cultures lacking astrocytes were generated from E17 rat cortex and after 14 days *in vitro* (DIV) they were infected with recombinant Herpes Simplex Virus (HSV) engineered to express WT or G86R mutant SOD1 or WT or M337V mutant TDP43. Rationale for utilizing this experimental platform is detailed in the STAR methods. By phase contrast microscopy, our cultures displayed abundant healthy-appearing neurons with a rich investment of neurites in the intercellular space (Figure 1A.). Immunocytochemically, a mesh of MAP2(+) dendrites were decorated with Piccolo-positive pre-synaptic terminals (Figure 1B.). Puncta of glutamate receptor subunits Gria1 and Gria2 opposing Piccolo-positive pre-synaptic terminals were abundant as well (Supplementary Figure 1). The co-localization of post synaptic density protein of 95 kDa molecular weight (PSD95) with Gria1 puncta as well as co-localization of Gria1 and GluN1 puncta re-enforce the notion that *bona fide* synaptic structures were assembled in our cultures (Supplementary Figure 1). By Western blot, our culture system displayed abundant expression of markers of excitatory and inhibitory neurotransmission and little evidence of astrocyte markers (Figure 1C.). These observations suggest that our experimental platform captures several network features seen *in vivo* – this is important because neuronal activity and metabolism are intimately linked (Dienel, 2019).

**Figure 1.**
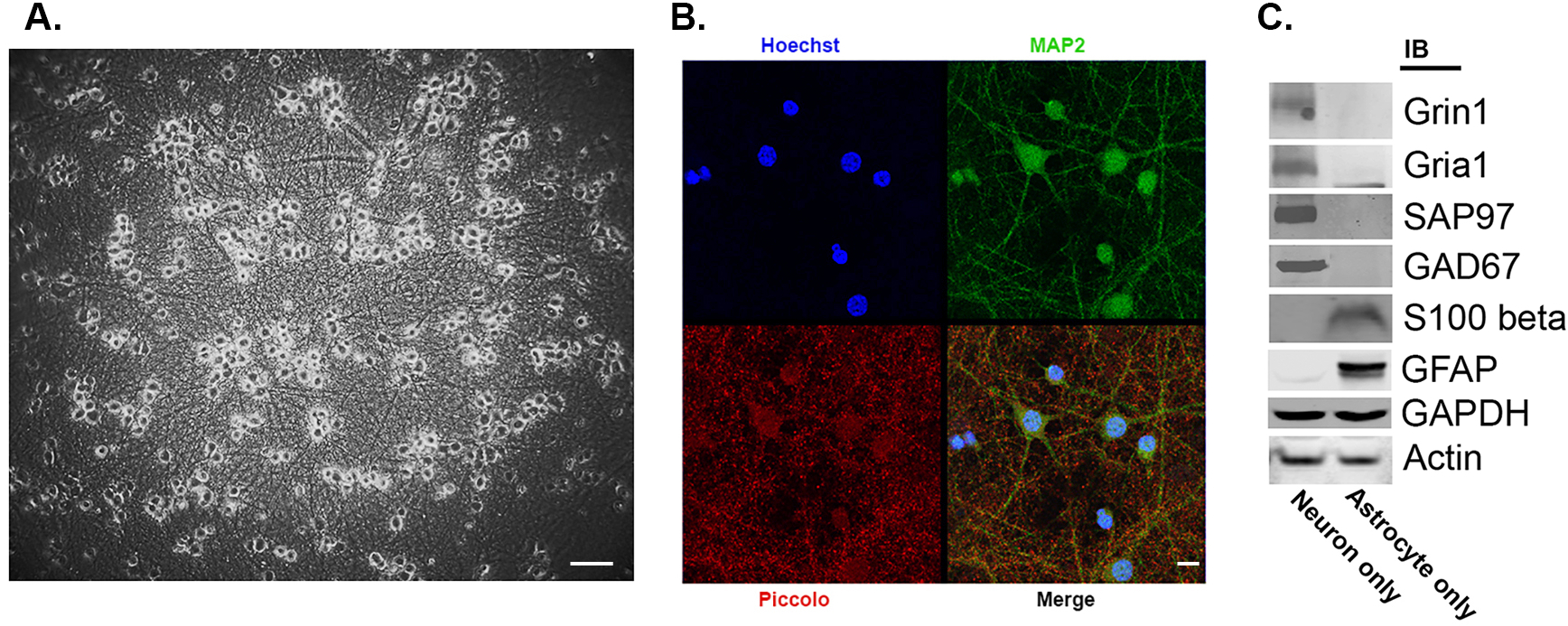
Characteristics of DIV 14 pure rat cortical neuron cultures. Panel A. Phase contrast image of cultures show abundant phase bright cell bodies with an extensive mat of interstitial neurites. Calibration bar = 85 microns. Panel B. A single confocal slice of cultures immunostained for MAP2 (green) and Piccolo (red); nuclei stained with Hoechst 33342. The merge of individual panels shows abundant pre-synaptic elements apposed to dendrites. Calibration bar = 25 microns Panel C. Western blot of cultures reveals evidence for excitatory (e.g., glutamate ionotropic receptor NMDA type subunit 1, Grin1; glutamate ionotropic receptor AMPA type subunit 1, Gria1; synapse associated protein of 97 kDa, SAP97) and inhibitor synapses (e.g., glutamate decarboxylase 67 kDa, GAD67) in cultures of neurons but not astrocytes (S100 calcium binding protein B, S100beta; glial fibrillary acidic protein, GFAP). Loading control was glyceraldehyde-3-phosphate dehydrogenase (GAPDH) when studying GFAP and actin for all the remaining proteins.

Using [1,2 ^13^C2]-D-glucose to track fuel utilization, we found that trends of metabolic alterations were similar for SOD1 and TDP43 expressing neurons in comparison with the control condition (neurons expressing LacZ). Net glucose uptake (i.e., CMRglc, the rate of glucose utilization) from the media (calculated by difference, glucose concentration before minus that after the 4 h incubation) by LacZ expressing neurons was about 3.2-3.8 μmol/mg/h, and net uptake was significantly reduced by about 25- 45% in WT SOD1, mutant SOD1 and mutant TDP43 expressing neurons (Figure 2 A., B.). This was associated with reduced lactate production and release to the medium in SOD1 and TDP43 expressing neurons, although the WT SOD1 group did not reach statistical significance (Figure 2 C., D.). For example extracellular lactate production in LacZ expressing neurons is approximately 1.2 μmol/mg/h while the lactate release from mutant SOD1 expressing neurons was approximately 0.80 μmol/mg/h and from mutant TDP43 expressing neurons approximately 0.68 μmol/mg/h. Isotope enrichment (atom percent excess, APE, see Star Methods) in lactate is reduced more in mutant SOD1 expressing neurons in comparison with WT SOD1 expressing neurons. In contrast, total lactate production and isotope enrichment in lactate were reduced more in WT TDP43 expressing neurons in comparison with mutant TDP43 (although both were lower the values in LacZ expressing neurons) (Figure 2 E., F.). Together these observations reveal a reduction in glucose utilization and glycolysis in the mutant SOD1 and mutant TDP43 expressing neurons (Figure 2, M. – P.).

**Figure 2.**
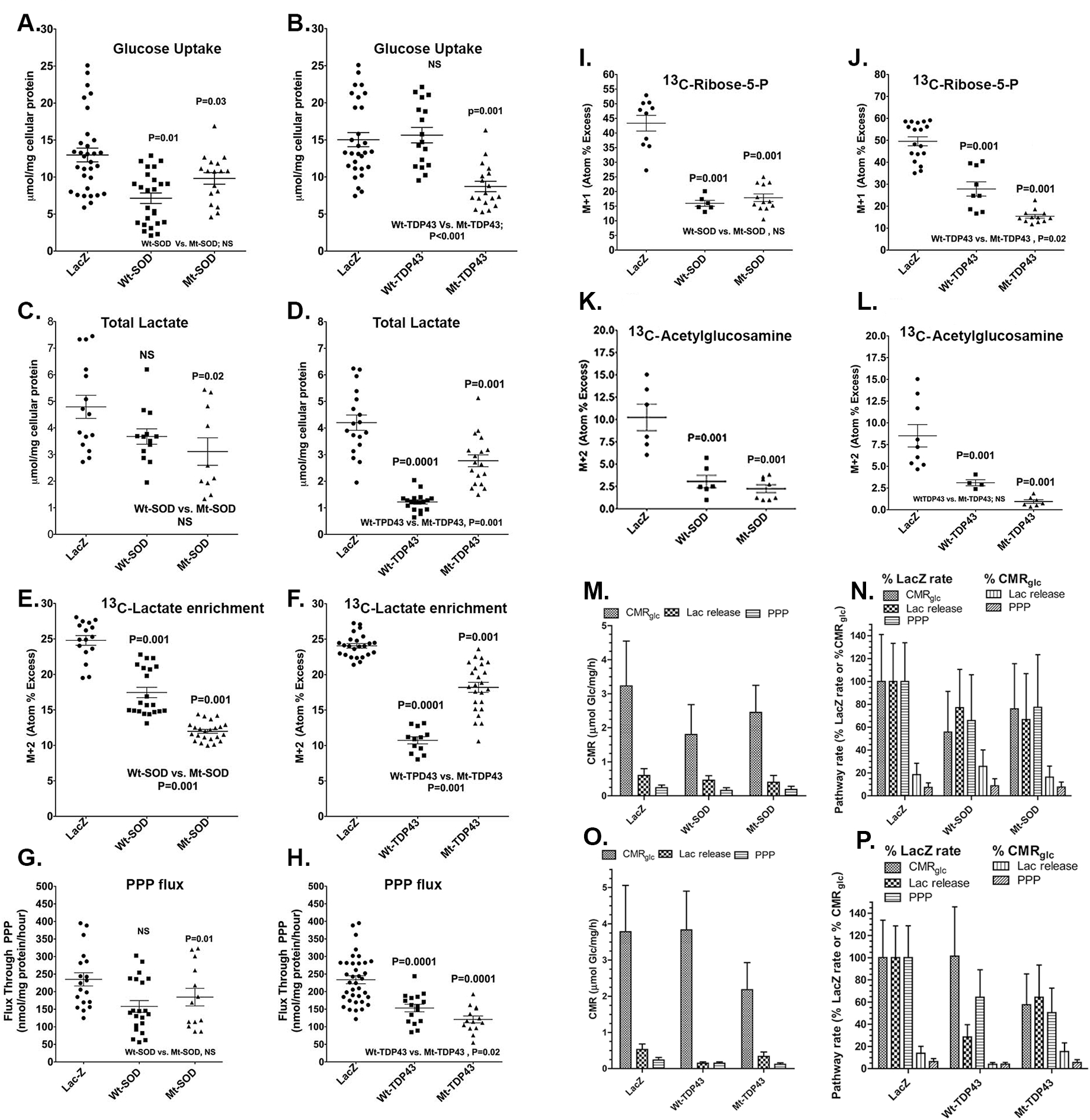
Metabolic interrogation of glycolysis and related pathways in neurons expressing WT and fALS genes. Pooled data from 3-5 experiments comparing neurons expressing LacZ with WT SOD1/TDP43 with mutant SOD1/TDP43 is presented. Horizontal lines denote means ± SD. ANOVA showed group differences and values above the scattergram indicate that in *post hoc* tests the specific group is statistically significantly different from the LacZ group (p value provided) or not different (NS - not significant). Values below the scattergram indicate that the WT or mutant proteins group are statistically significantly different or not. Panel labels: A, B, Net glucose uptake during the 4 h incubation; C, D, total lactate release to medium; E, F, ^13^C enrichment of medium lactate; G, H, Flux of glucose through the pentose phosphate pathway (PPP); I, J, ^13^C enrichment of intracellular ribose-5-phosphate (P); K, L, ^13^C enrichment of intracellular acetylglucosamine. For the CMRglc summary panels (M. and O.), data are means and 1 SD, with all rates in units of glucose or its equivalent (i.e., lactate/2). For the pathway rates expressed as percent of the LacZ rate or of CMRglc (Panels N. and P.), values are mean quotient and vertical bars are CVs for the 100% rates or SDs that account for error propagation for all others. Statistically significant changes are indicated in the panels that illustrate the primary data.

Beyond its role in producing ATP and pyruvate (for entrance via acetyl CoA into the TCA cycle) the metabolism of glucose by glycolysis generates intermediates that serve other cellular functions. The glycolytic intermediate glucose-6-phosphate [Glc-6-P] enters the PPP. The PPP rate is 0.24 μmol/mg/h in LacZ expressing neurons and 0.19 μmol/mg/h for mutant SOD1 and 0.12 μmol/mg/h for mutant TDP43 expressing neurons (Figure 2 G., H.). When expressed as a fraction of CMRglc, the magnitude of the relative PPP rate (i.e., about 7-9% of CMRglc in LacZ expressing neurons) fell in proportion to CMRglc in SOD1 neurons, whereas it fell from 6.3% in LacZ neurons to 5.6% in mutant TDP43 expressing neurons (Figure 2 N. – P.). These lower PPP rates were associated with reductions in ribose-5-phosphate levels (a product of PPP) that were disproportionately greater than the respective fall in PPP rate in the mutant SOD1 and TDP43 expressing neurons (Figure 2 I., J.). The glycolytic intermediate fructose-6- phosphate enters into the HBS pathway. Monitoring acetylglucosamine levels reports on this pathway, and the isotope enrichment in acetylglucosamine in SOD1 and TDP43 expressing neurons was dramatically reduced, more than the fall in glycolytic rate, PPP rate, and ribose-5-phosphate enrichment (Figure 2 K., L.) By several measures neurons expressing mutant proteins have more severe deficits that neurons expressing WT proteins. Net glucose uptake, medium lactate levels, PPP flux, and ribose-5-phosphate enrichment did not differ in the mutant and WT SOD1 expressing neurons but isotope enrichment in lactate was lower in mutant SOD1 vs. WT SOD1 expressing neurons. Net glucose uptake, PPP flux, and ribose-5-phosphate enrichments were lower in mutant TDP43 vs. WT TDP43 expressing neurons. We provide a summary of CMRglc and CMRglc for SOD1 and TDP43 expressing neurons as a percentage of LacZ neurons (Figures 2 N and P respectively.).

The fall in glycolysis in SOD1 and TDP43 expressing neurons could impact the operation of the TCA cycle and so next we looked at the labeling of TCA cycle intermediates by [^13^C]Glucose. Isotope enrichment in citrate, fumarate and malate is maintained or even increased in neurons expressing SOD1 and TDP43 (Figure 3A. – F.). There is statistically significantly more isotope enrichment in fumarate and in malate in mutant SOD1 or TDP43 compared with the WT versions of these proteins. These observations imply that the entrance of pyruvate into the TCA cycle is maintained or enhanced despite the mutant protein-evoked reduction in glycolytic net labeling of glucose.

**Figure 3.**
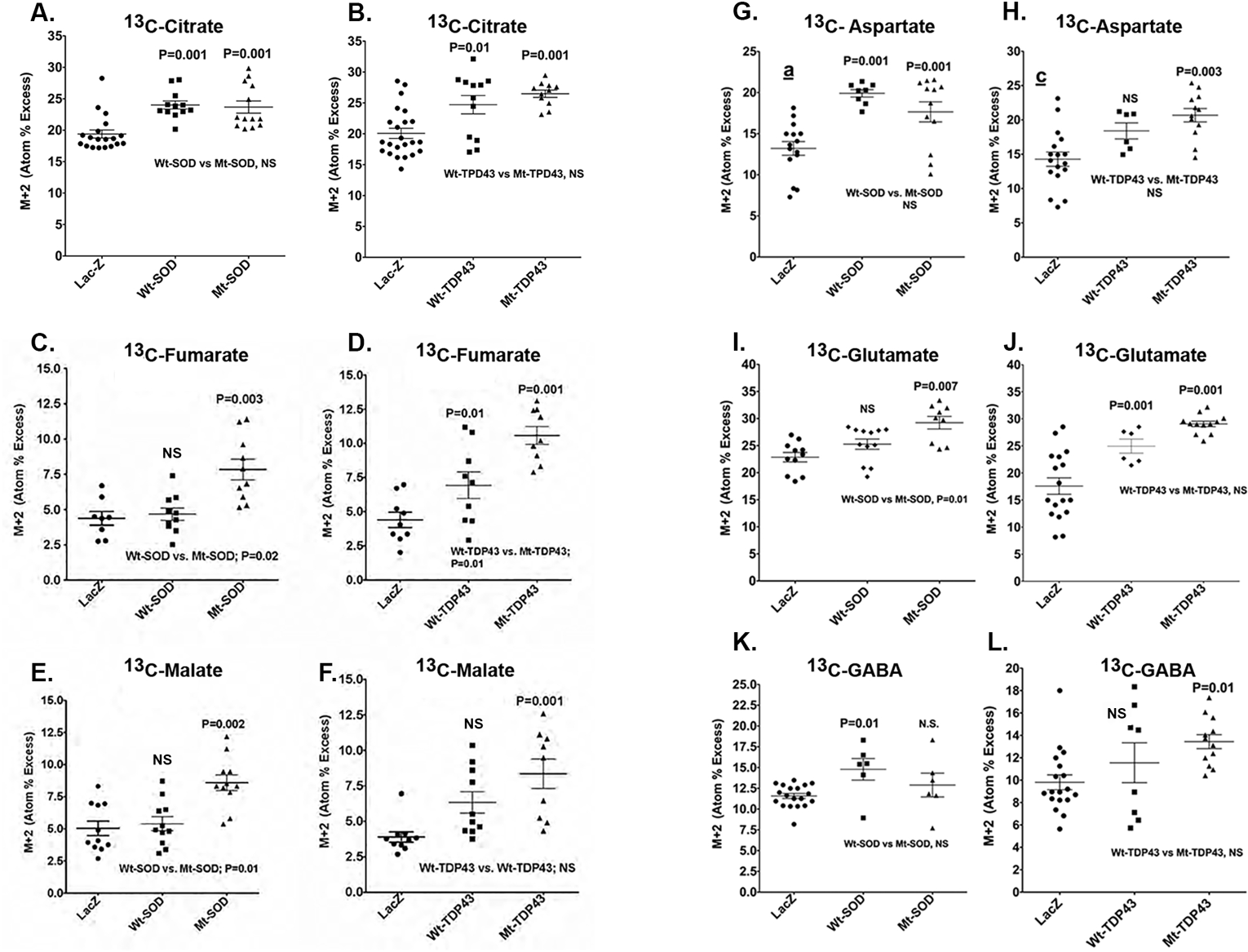
Metabolic interrogation of TCA cycle and related pathways in neurons expressing WT and fALS genes. Pooled data from 3-5 experiments comparing neurons expressing LacZ with WT SOD1/TDP43 with mutant SOD1/TDP43 is presented. Horizontal lines denote means ± SD. ANOVA showed group differences and values above the scattergram indicate that in *post hoc* tests the specific group is statistically significantly different from the LacZ group (p value provided) or not different (NS - not significant). Values below the scattergram indicate that the WT or mutant proteins group are statistically significantly different or not. ^13^C enrichment of tricarboxylic acid (TCA) cycle constituents (A-F) and TCA cycle-derived amino acids (G-L).

In addition to TCA cycle intermediates we monitored major amino acids derived from the TCA cycle, and we find significantly increased isotope enrichment in glutamate and aspartate in mutant SOD1 and mutant TDP43 expressing neurons when compared with LacZ expressing neurons (Figure 3G. – J.). There is statistically significantly higher isotope enrichment in glutamate in mutant SOD1 but not the mutant TDP43 compared with the WT versions of these proteins. There is significantly increased isotope enrichment in GABA in WT SOD1 neurons as well as mutant TDP43 neurons (Figure 3L.). We also measured total amino acid levels and found virtually no differences between the abundance of 19 amino acids in neurons expressing the WT or mutant proteins, although small group differences were seen when comparing LacZ to WT or mutant SOD (Supplemental Table 1). The enhanced labeling of the TCA cycle-derived amino acids by ^13^C-glucose, indicates that there must be additional source(s) for the four-carbon unit for the synthesis of glutamate, GABA, and aspartate. The requirement for a carbon source arises from the lack of pyruvate carboxylase in neurons, which renders them incapable of synthesizing the 4-carbon precursor, oxaloacetate, *de novo* from glucose. In brain, this process is carried out mainly by astrocytes and to a much lesser extent oligodendroglia. In our experimental platform a likely source of four-carbon units is growth media amino acids. Note that the additive Glutamax contains a dipeptide L-alanyl-L-glutamine, a precursor of glutamate, GABA and aspartate.

The alterations in glycolysis and TCA cycle evoked by mutant SOD1 and mutant TDP43 expressing neurons (in comparison with LacZ expressing neurons) might be expected to impact cellular energy levels and redox status. We find there are remarkably minimal effects on the concentrations of adenine nucleotides or the levels of NAD^+^ and NADH between experimental groups (Supplemental Table 2). ATP and AMP levels were significantly increased in mutant SOD1 expressing neurons and ATP was also elevated in WT TDP43 expressing neurons. We find a statistically significant reduction in NADH and an increased in NADPH in WT TDP43 expressing neurons (in comparison with LacZ) as well as reductions in NAD^+^ and NADP^+^ in WT SOD1 expressing neurons. Also some groups showed differences between SOD1 or TDP43 expression and LacZ. If these changes in NADP+ or NADPH levels were a reflection of oxidative stress, we would anticipate corresponding alteration in the ratio of reduced to oxidized glutathione (e.g., the GSH/GSSG ratio.) In fact, we see that expression of WT or mutant SOD1 or TDP43 has essentially no effects on GSH or GSSG levels. This argues against a significant alteration in redox status under our experimental conditions. It is possible that fuel utilization by mutant protein-expressing neurons is altered is a way that maintains neuronal redox status and energy charge.

In light of the above described alterations in fuel utilization we wondered about activation of AMP dependent protein kinase (AMPK), a master sensor and regulator of intermediary metabolism. In comparison with LacZ expressing neurons we found neither alterations in AMPK activation nor its downstream targets acetyl-CoA carboxylase (ACC) and factor 4E binding protein (4EBP1) in WT or mutant SOD/TDP43 expressing neurons (Supplemental Figure 2). Thus, while expression of fALS-causing proteins re-wire metabolism, the production of ATP is sufficient to obviate an AMPK-dependent adjustment of anabolism to catabolism.

In sum, expression of ALS-causing mutant proteins SOD1 and TDP43 in cortical neurons leads to reduced glycolytic, PPP and HBS pathway net labeling by glucose but this is unassociated with deficits in energy and redox status. The TCA cycle operates at a higher than normal level with glucose-derived carbon skeletons entering as pyruvate and likely oxidized medium amino acids resulting in maintenance of ATP levels (summarized in Supplemental Figure 3). While some measures of mitochondrial physiology deviate from normal in ALS (Mattiazzi et al., 2002) (Bowling et al., 1993; Dupuis et al., 2004; Jung et al., 2002; Magrané et al., 2012), three reporters of mitochondrial operations in our neuronal models are remarkably normal: 1) ATP/ADP ratio, 2) GSSG/GSH ratio and 3) the abundance of TCA cycle intermediates such as aspartate. This argues that altered mitochondrial function in our models is not observed.

### Functional consequences of glycolysis on ALS causing mutant proteins

While we have described alterations in intermediary metabolism evoked by disease causing proteins, we do not know if such rewiring that reduces glucose utilization has consequences for neuronal function and survival. We hypothesized that studying the effects of ablation of individual genes in metabolic pathways (beginning with glycolysis) would be a productive approach to address this rewiring issue (see STAR methods). To pursue this in an expeditious manner we undertook a growth fitness assay in *S. cerevisiae* engineered to inducibly express wild type or Q331K, mutant TDP43 in a wild type background or in the presence of deletions of different glycolysis-associated genes (Supplemental Table 3.). Non-essential genes (e.g., not absolutely required for survival) were studied in a haploid yeast strain with a deletion of the corresponding single allele. Essential genes (e.g., *PGI1, GFA1, FBA1, TPI1, PGK1, GPM1* and *PYK1*) were studied in a diploid yeast strain with a deletion of one of the two alleles. We developed a semi-quantitative assay evaluation that took into account three variables affecting cellular growth: 1) the expression of the transgene (Wt TDP-43 or Mt TDP-43), 2) the presence or absence of the gene of interest in glycolysis, and 3) the use of different carbon fuels necessary for either expressing the transgene (galactose, Gal) or inhibiting transgene expression (glucose, Glc) (Figure 4 A.). The crosstalk between the first and second variables on cellular growth reflect the main signal and the third variable is a source of noise and its magnitude may depend on variable two. Equation 1 reflects the effect of expressing TDP-43 on cellular growth, in the absence of a specific metabolic gene-of-interest. The additional effect of using different carbon fuels (glucose/galactose) is factored in. Equation 2 reflects the effect of expressing TDP-43 on cellular growth, in the presence of the specific metabolic gene-of-interest. Also here, the additional effect of using different carbon fuels (glucose/galactose) is factored in. The ratio of equation #1 to #2 (equation #3) reflects the specific effects of ablating a specific gene-of-interest on TDP-43 impairment of growth fitness. To validate our approach, we examined the effects of a gene previously found in a yeast screen as a suppressor of TDP43 growth fitness (Armakola et al. 2012) (Figure 4B.). In our assay, deletion of this gene, *dbr1,* promoted growth fitness confirming that our semi-quantitative assay system can be deployed to find modifiers of toxicity.

**Figure 4:**
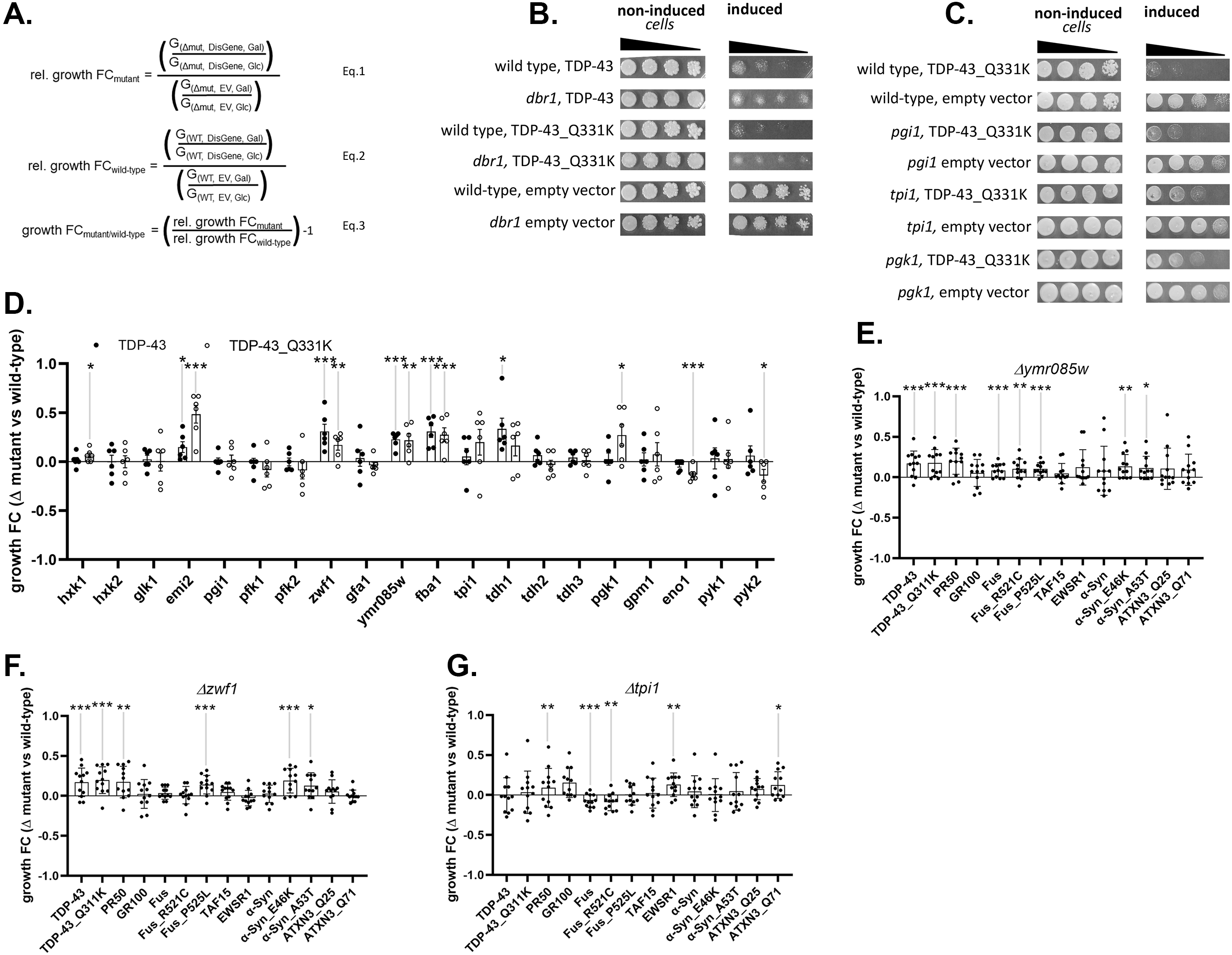
Systematic deletion of individual glycolytic enzymes reveal modifiers of protein toxicity of neurodegenerative disease-associated proteins in yeast. Panel A. Equations used in semi-quantitative growth fitness assay take into account three variables: 1) the presence or absence of a specific metabolism gene (noted as WT or Δmut), 2) the presence of the empty Expression Vector or Disease Gene elaborated by the expression vector (noted as EV or DisGene), and 3) growth in Galactose or Glucose (noted as Gal or Glc). Note, the DisGene is expressed under the control of a galactose inducible promoter, and DisGene expression is inhibited in the presence of glucose. Equation one (Eq.1) describes the relative growth fold changes (“rel. growth FC”) in the yeast strain mutant (e.g. lacking) a metabolism gene, expressing the disease gene or not and growing on galactose or glucose. Equation two (Eq.2) describes the rel. growth FC of yeast strain WT for the metabolism gene, expressing the disease gene or not and growing on galactose or glucose. Equation three (Eq. 3) describes the specific effect of the metabolism gene on DisGene growth fitness by dividing Eq.1 by Eq2. yielding the growth FCmutant/wild-type. Panel B. As previously published (Armakola et al., 2012), deleting *dbr1* partly rescues Wt and Mt (Q331K) TDP43-induced protein toxicity in the corresponding yeast dilution models. Panel C. Representative examples of glycolytic enzyme deletion strains tested for effects on TDP43 toxicity. The strain deleted for *pgi1* did not affect protein toxicity, whereas strains deleted for *tpi1* or *pgk1* reduced Mt TDP- 43 toxicity. Panel D. Screening deletion strains for glycolytic enzymes and for branching enzymes into PPP and HBS pathways using the Wt (filled circles) and Mt (open circles) TDP-43 protein toxicity models; n=6. Growth fitness benefits (group FC > 0) were seen in yeast deleted for *hxk1, emi2, zwf1, ymr085w, fba1 tdh1* and *pgk1*. Growth fitness decrement (growth FC < 0) were seen in yeast deleted for *eno1* and *pyk2.* Panel E. Growth fitness benefits were seen in yeast deleted for *ymr085w* on a variety of proteotoxic insults relevant to ALS (e.g., TDP43, C9orf72 DPRs, FUS, TAF15, EWSR1) Parkinson’s Disease (e.g., α-Syn) and Spinocerebellar Ataxia (e.g., ATXN3); n=12. Growth fitness benefits were seen in yeast expressing Wt and Mt TDP43, PR50, FUS (Wt and P525L) and Wt and Mt (E46K and A53T) α-syn. Panel F. Growth fitness effects of ablation of *zwf1* on a variety of proteotoxic insults relevant to ALS, PD and SCA; n=12. Growth fitness benefits were seen in yeast expressing Wt and Mt TDP43, PR50, Mt FUS and Mt (E46K and A53T) α-syn. Panel G. Growth fitness effects of ablation of *tpi1* on a variety of proteotoxic insults relevant to ALS, PD and SCA; n=12. Growth fitness benefits were seen in yeast expressing PR50, Wt and Mt FUS, EWSR1 and ATXN3 Q71.(*p<0.05, **p<0.01, ***p<0.005).

Using this assay platform we assayed 20 non-essential glycolysis associated genes found that deletion of 5 genes enhanced growth fitness of *S. cerevisiae* upon induction of mutant TDP43 expression: *emi2 (*paralog of yeast glucokinase)*, zwf1 (*Glc-6-P dehydrogenase)*, ymr085w (*paralog of yeast glutamine-fructose-6-phosphate amidotransferase)*, fba1* (fructose 1,6 bisphosphate aldolase), and *pgk1 (3- phosphoglycerate kinase)* – with enzyme names of mammalian homologs or orthlogs in parentheses, respectively (Figure 4C., D.). In 3/5 cases, the deletion also improved the growth fitness upon induction of wild-type TDP43 expression. In contrast, we found that deletion of 2 genes reduced growth fitness of *S. cerevisiae* upon induction of mutant (but not wild-type) TDP43 expression: *eno1* (enolase) and *pyk2* (pyruvate kinase).

Next we asked if deletion of the five genes of interest (e.g., *emi2, zwf1, ymr085w, fba1* or *pgk1*) improve the growth fitness of additional yeast-based models of familial ALS or other neurodegenerative diseases. To address this issue, we studied yeast that inducibly-express two different toxic diamino acid peptides (DPRs) derived from the *C9ORF72* mutation, wild-type or R521C or P525L fused-in-sarcoma (FUS), wild type TAF15, wild type EWSR1, wild type or E46K or A53T alpha synuclein, or ATXN3 with 25 glutamines (Q25) or 71 glutamines (Q71). We found that *ymr085w* and *zwf1* had broad growth fitness phenotypes, showing beneficial actions in 8 and 6 different models, respectively (Figure 4E., F.). In contrast, *tp1i* which did not show a beneficial effect in the original TDP43 screen, had mixed effects (Figure 4 G.) while *emi2, fba1, tdh1* and *pgk1* had no consistent effects in these models (Supplemental Figure 4) In sum, we find that ablation of several yeast genes involved in glycolysis confer growth fitness benefits on TDP43 based models of ALS and at least two genes (*ymr085w* and *zwf1)* confer growth fitness benefits in yeast-based models of several different models of ALS and neurodegeneration in general. *zwf1* (G6PDH) is the first step in PPP and *ymr085w* (GF6PA) is the first step in the HBS pathway. An implication of removal of these enzymes is that glucose-derived carbon cannot enter the respective branch pathways, thereby altering unidentified processes (e.g., glucose flux into different catabolic routes, actions of regulatory molecules, or unanticipated effects of metabolic shifts) that have favorable outcomes.

### Effects of reducing G6PDH in a C.elegans model of TDP43 proteinopathy

The yeast growth fitness platform is a powerful tool for finding modifiers of mutant gene phenotypes. The extent to which a pleiotropic phenotype like growth fitness reports on neuronal biology requires secondary screens. One experimental platform we next deployed was a *C.elegans* model of TDP43 proteinopathy. The advantages of this model are that: 1) it is an *in vivo* system with an intact nervous system, and 2) a proven track record of bridging cellular models of biology to mammalian systems (Jablonski et al., 2015; Lim et al., 2012; Periz et al., 2015; Zhai et al., 2015). We used feeding RNAi to knockdown genes of interest in two different strains (Supplemental table 3.).

CL6303 worms have nervous system specific expression of human WT TDP43 and are engineered to have enhanced neuronal RNAi. RK179 worms do not express human TDP43 but have the identical genetic enhancements of neuronal RNAi. After two generations of growth on RNAi bacteria (targeting *gspd-1, gfat-1, gfat-2 or tdp-1* – the worm orthologs of G6PDH, GF6PA and TDP43), we studied various locomotor phenotypes. We find that RNAi targeting *gspd-1* confers a locomotor benefit specifically to the human TDP43 expressing animals (Figure 5). The beneficial effects on locomotion of reducing G6PDH on TDP43 phenotypes was an encouraging confirmation manipulations of intermediary metabolism are worthy of study.

**Figure 5.**
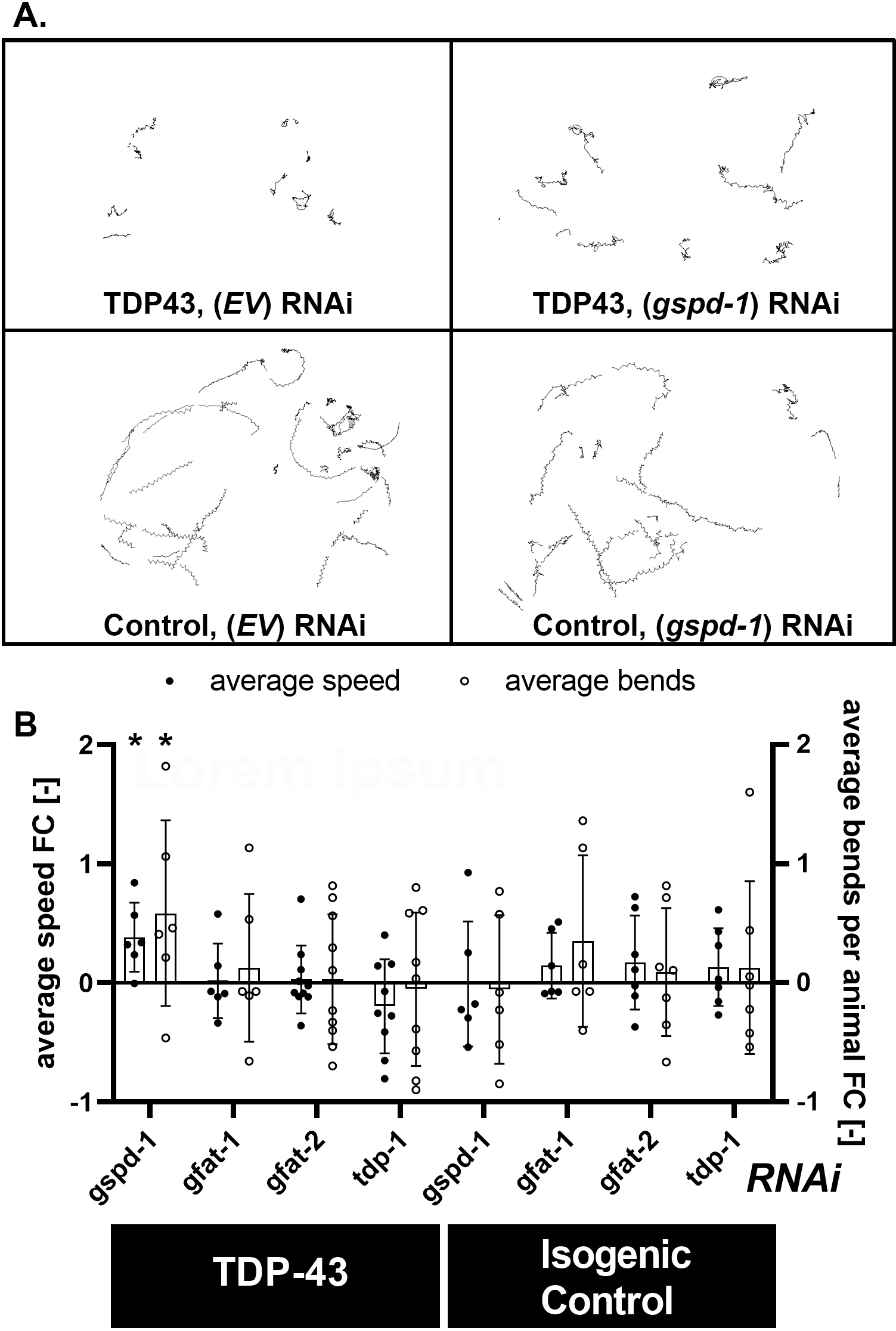
Knock down of worm G6PDH (*gspd-1*) suppresses the locomotor defect incurred by nervous system expression of Wt human TDP43. Panel A. Exemplary displacement paths acquired from swimming worms in M9 medium recorded with a high-resolution camera using the ImageJ wrMTrck plugin. Paths were collected from animals expressing TDP43 (CL6303) and isogenic controls (RK179); feeding RNAi targeting *gspd-1* or vector controls (EV). Panel B. Quantitative analysis of average speed (pixels/minute, filled circles) and average beats per animal (1/minute, open circles) reveals that average speed and bend rates of TDP43 expressing animals are improved by knock down of *gspd-1* but not knock down *of gfat-1, gfat-2 or tdp-*1 (in comparison with EV control) . There is no effect of RNAi knockdown of any of the studied genes in isogenic control animals that do not express TDP43. Thus the beneficial effect of reducing *gspd-1* is specific to animals expressing TDP43. Effects were quantified as average animal speeds and bending rates of 9-15 L4 animals per run; n=6. (**p<0.01).

### Effects of reducing G6PDH in mammalian neuron models of ALS

Based on the promising results described above, we asked if manipulation of G6PDH in a primary rat neuron model system of TDP43 proteinopathy had beneficial effects. We began by generating a series of microRNAs targeting the rat *G6PDH* and selected one that achieves a 65% knockdown of the protein (Figure 6A.). Next we generated embryonic rat spinal cord cultures grown on astrocytes and after 14 days *in vitro* (DIV) they were infected with Herpes Simplex Virus engineered to express LacZ, wild-type human TDP43 or human M337V TDP43 ± the miRNA targeting *g6pdh* or a scrambled targeting sequence (scr) miRNA. In prior work we showed that infection of cultures with 2 recombinant viruses achieves a >90% co-infection rate of neurons (Mojsilovic-Petrovic et al., 2009). When we counted motor neurons 5 days later, we find that M337V TDP43 kills ∼ 50% of motor neurons and knockdown of G6PDH leads to a statistically significant blunting of this noxious effect whereas G6PDH knockdown had no adverse effect on the survival of LacZ or wild type TDP43 expressing neurons (Figure 6B., C).

**Figure 6.**
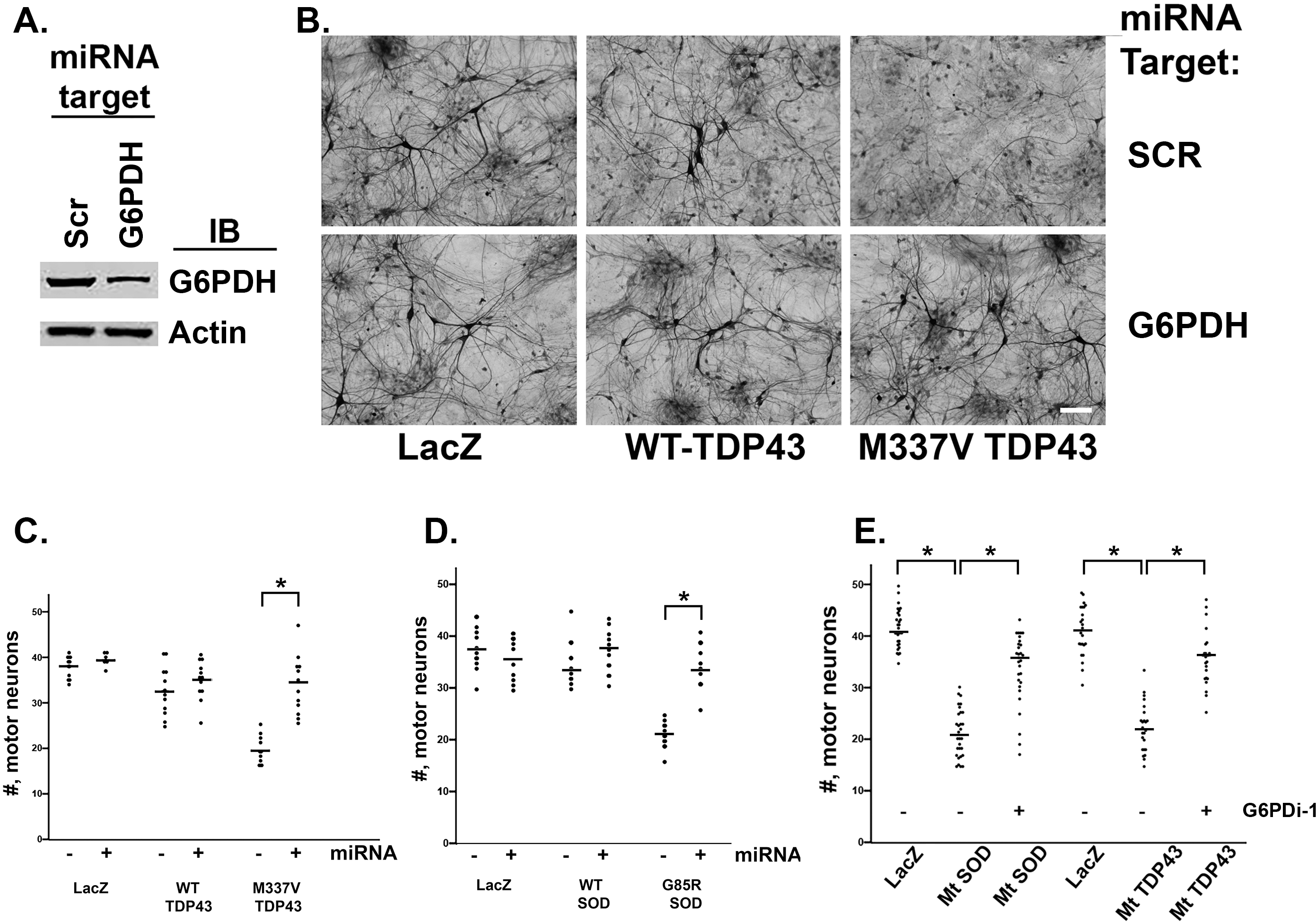
Reduction of G6PDH abundance or activity protects against the motor neuron toxicity of mutant SOD1 and mutant TDP43. Panel A. Lysates from pure neuronal cultures infected with HSV engineered to express a miRNA targeting G6PDH or a scramble (scr) sequence miRNA were immunoblotted for G6PDH and actin. The miRNA targeting G6PDH reduces neuronal G6PDH expression by ∼60% of the control. Panel B. Representative photomicrographs of SMI32 immunostained DIV14 spinal cord cultures expressing LacZ, WT-TDP43 or M337V mutant TDP43 and either the miRNA targeting G6PDH or the scramble sequence miRNA. Neurons with cell bodies larger than 25 microns are motor neurons. Fewer motor neurons are evident in the M337V mutant TDP43 + miRNA scr in comparison with the M337V mutant TDP43 + miRNA G6PDH. Scale bar = 75 microns. Panel C. There is no statistical difference in the number (denoted by #) of motor neurons expressing LacZ with or with knockdown of G6PDH. Similarly, expression of Wt TDP43 does not lead to motor neuron death with or without knockdown of G6PDH. Expression of M337V Mt TDP43 does lead to a significant loss of motor neurons and knockdown of G6PDH leads to a statistically significant blunting of the Mt TDP43 toxicity. Results from ≥ 4 experiments, *, p < 0.05. Panel D. There is no statistical difference in the number of motor neurons expressing LacZ with or with knockdown of G6PDH. Similarly, expression of Wt SOD1 does not lead to motor neuron death with or without knockdown of G6PDH. Expression of G85R Mt SOD1 does lead to a significant loss of motor neurons and knockdown of G6PDH leads to a statistically significant blunting of the Mt SOD1 toxicity. Results from ≥ 4 experiments, *, p < 0.05. Panel E. A three way comparison of number of motor neurons expressing LacZ, Mt SOD1 – G6PDi-1 and Mt SOD1 + G6PDi-1. Statistically significant group differences were seen by ANOVA and post hoc test reveal that Mt SOD1 – G6PDi-1 is significant different from LacZ and from Mt SOD1 + G6PDi-1. Results from ≥ 4 experiments, *, p < 0.05. A three way comparison of number of motor neurons expressing LacZ, Mt TDP43 – G6PDi-1 and Mt TDP43 + G6PDi-1. Statistically significant group differences were seen by ANOVA and post hoc test reveal that Mt TDP43 – G6PDi-1 is significant different from LacZ and from Mt TDP43 + G6PDi-1. Results from ≥ 4 experiments, *, p < 0.05.

Encouraged by these observations and noting the similarity of metabolic rewiring evoked by mutant forms of TDP43 and SOD1, we studied the effects of G6PDH reduction in cultures expressing wild type SOD1 or G85R SOD1. We find that G85R SOD1 kills ∼ 50% of motor neurons and knockdown of G6PDH leads to a statistically significant blunting of this noxious effect (Figure 6D.). Again, G6PDH knockdown had no adverse effects on LacZ or wild type SOD1 expressing neurons.

Finally we looked at two other fALS models involving the expression of GA50 or PR50, noxious DPRs that are generated from the hexanucleotide expanded mutant version of *C9ORF72*(Ash et al., 2013; Gupta et al., 2017; Mizielinska et al., 2014; Wen et al., 2014). We find that GA50 or PR50 kills ∼ 50% of motor neurons and knockdown of G6PDH leads to a statistically significant blunting of this noxious effect (Supplemental Figure 5).

The primary goal of research into neurodegenerative diseases is to find targets for manipulation by small molecules, i.e., drugs. In light of our observations on the contribution of G6PDH to neuronal dysfunction in ALS models, we wondered whether this was a druggable target. Several different G6PDH inhibitors have been described but their specificity is open to question. A new highly, selective small molecule G6PDH inhibitor was reported last year, “G6PDi-1” (Ghergurovich et al., 2020), and we tested it in our system. We find that G6PDi-1 has no effects on the survival of motor neurons expressing no transgene, LacZ or WT SOD1 or WT TDP43. Motor neuron death evoked by the expression of mutant SOD1 or mutant TDP43 is robustly suppressed by administration of G6PDi-1 (Figure 6E.).

Together these results suggest that reducing the abundance or inhibiting the activity of G6PDH in neurons confers a broad beneficial effect on cellular phenotypes evoked by expression of fALS genes.

### Metabolomic effects of G6PDH knockdown in neurons expressing fALS proteins

The conversion of Glc-6-P to 6-phosphogluconolactone by G6PDH is the first and rate-limiting enzyme in PPP. The ratio of Glc-6-P that enters glycolysis (via metabolism to fructose-6-phosphate) to PPP is ∼ 10:1 and we considered the possibility that reducing G6PDH levels could make this ratio even larger. If this were true, it might enhance glycolytic flux and potentially help mend the demonstrated glycolytic flux defect we find in fALS expressing neurons (Figure 2). To examine this issue, we undertook glucose isotopologue interrogations of pure neuron cultures expressing fALS genes ± 65% knockdown of G6PDH that blunts motor neuron death (Supplemental Figure 6). In comparison with a scrambled sequence miRNA, the miRNA targeting G6PDH had no effects on glucose uptake, total lactate, isotope enrichment in lactate, flux through PPP, isotope enrichment in R-5-P or isotope enrichment in acetylglucosamine in neurons expressing mutant SOD1 or mutant TDP43. In terms of TCA cycle intermediates, reduction of G6PDH on motor neuron survival led to increased isotope enrichment in malate only in mutant TDP43, but not SOD1, expressing neurons. In terms of amino acids, reduction of G6PDH reduced led to isotope enrichment in aspartate and glutamate only in the mutant SOD1 expressing neurons. Together these observations lead to two conclusions: 1) knockdown of G6PDH has no effect of the flux of glucose through glycolysis in neurons expressing fALS proteins, and 2) the beneficial effects of manipulation of G6PDH in the presence of mSOD1 or mTDP43 are metabolically distinct and thus potentially mechanistically distinct.

## Discussion

Here we have investigated the intermediary metabolism of neurons that express fALS genes. We find that neurons expressing two different fALS mutant proteins display a reduction of net utilization of glucose via glycolysis, PPP and HBS pathways. At the same time the fuel entrance into TCA cycle is maintained or increased. This is may be due to enhanced entry of pyruvate and likely, as well as, carbon skeletons from growth medium amino acids or glutamine. While mitochondrial dysfunction has been described in some ALS disease models, we find that in neurons expressing fALS genes mitochondrial operations are sufficiently robust to maintain ATP levels and healthy redox status. It appears that fALS genes provoke quantitative (and potentially qualitative) alterations to metabolic pathways potentially as a mechanism to maintain ATP and redox status. The case that this is pathophysiologically significant derives from the observation that loss-of-function genetic maneuvers at specific steps of the glycolysis pathway suppress toxic phenotypes. Null or partial reduction in G6PDH has broad beneficial effect in three experimental platforms (e.g., *S. cerevisiae, C. elegans* and rodent spinal cord neurons). This is unlikely to be due to a specific effect on glycolysis and the mechanisms by which reducing G6PDH blunts the toxicity of mSOD1 versus mTDP43 may be distinct.

Alterations in the energy economy in fALS-based experimental models has been recognized for many years. Mitochondrial abnormalities (such as sequestration of toxic proteins and perturbed morphology and function) have been described in SOD1, TDP43, VCP, CHCHD10 and C9ORF72 models of ALS (Anderson et al., 2019; Baek et al., 2021; Bartolome et al., 2013; Chang et al., 2011; Choi et al., 2019; Israelson et al., 2010; Kim et al., 2013; Li et al., 2020a; Mattiazzi et al., 2002; Parone et al., 2013; Pasinelli et al., 2004; Straub et al., 2018; Tan et al., 2013; Wang et al., 2019; 2016; Yu et al., 2020). This is associated with alterations in glycolysis (Allen et al., 2014; Manzo et al., 2019; Valbuena et al., 2016), glutaminolysis (Valbuena et al., 2016), PPP (Allen et al., 2014; Manzo et al., 2019; Tefera et al., 2019), amino acid depletion (Tefera and Borges, 2019; Valbuena et al., 2016) and impaired glucose utilization (Manzo et al., 2019). The above noted observations are derived from non-neuronal cells or tissues composed of multiple cells types that differ in how they break down fuel to generate ATP and thus are silent on the issue of neuronal metabolism. We provide an in depth and quantitative analysis of fuel utilization, redox status and ATP/ADP/AMP abundance in a population of interconnected vertebrate neurons. We are open to the possibility that alterations in the metabolism of non-neuronal cells occurs co-incidentally as proposed by the work on oligodendrocytes (Lee et al., 2012) and astrocytes (Ferraiuolo et al., 2011) in disease models. Since glucose is the prime fuel for neuronal ATP production, our work highlights the fundamental alterations in neuronal intermediary metabolism incurred by fALS genes.

We find a consistent reduction in net labeling of glycolytic intermediates by [1,2 ^13^C2]-D-glucose in neurons expressing mSOD1 or mTDP43, an observation which complements work by the Borges and Zarensca laboratories in multi-cell type fALS models (Tefera et al., 2019), (Tefera and Borges, 2019) (Manzo et al., 2019). Glycolysis is regulated at a number of steps – classically, glucose uptake from the extracellular milieu and allosteric regulation of enzymes that engage in essentially irreversible reactions (e.g., hexokinase, phosphofructokinase and pyruvate kinase). Allosterically active small molecules include ATP, AMP, cellular pH, fructose 2,6 biphosphate (F-2,6-BP), F-1,6- BP, and alanine. In addition, glycolysis responds to mechanical extracellular cues, in particular the structure of the actomyosin cytoskeleton (Park et al., 2020). At present we do not know which of these points of control account for our finding of reduced glycolytic flux in neurons expressing mSOD1 or mTDP43.

One simple hypothesis is that reduction in glycolytic flux in neurons expressing mSOD1 or mTDP43 is the primary pathophysiological driver. If so, neurons might enact metabolic re-wiring to maintain ATP production and several observations are consistent with this notion. First substrate flux through PPP and HBS pathways are reduced and the TCA cycle operates normally or even at supra-normal levels. Our data suggest enhanced utilization of pyruvate and other sources of carbon skeleton, most likely growth medium amino acids. Second, the expression of mSOD1 or mTDP43 in neurons has minimal effects on adenylate nucleotide levels and redox status. Several observations, however, suggest that the simple hypothesis is incomplete. First, in our yeast screen we found that ablation of multiple, specific glycolytic enzymes conferred benefits in the growth fitness assay and this would be expected to reduce glycolytic flux. Second, knockdown of G6PDH was beneficial in multiple additional experimental platforms but does not, in fact, affect glycolytic flux. A simple explanation is that the beneficial metabolic consequences of ablation or knockdown of specific glycolysis-associated enzymes occurs by routing substrates for utilization into pathways we are not monitoring. We suspect, however, that this is not the complete story.

Our current model does not include consideration of glycolytic enzyme spatial distribution. For example, glycolytic enzymes such as glyceraldehyde 3-phosphate dehydrogenase (GAPDH) and phosphoglycerate kinase are located on transport vesicles and their local, “on-board” ATP production is required for fast axonal transport (Zala et al., 2013). Additionally, glycolytic enzymes have the capacity to undergo phase separation to form membraneless condensates that enhance substrate flux (Jin et al., 2017). Beyond subcellular localization, metabolic enzymes can have cell biological functions beyond catabolism. GAPDH interacts with ADP-ribosylation factor 1 GTP’ase activating protein to regulate coat protein I-mediated vesicle fission and this occurs independent of GAPDH’s catalytic activity (Yang et al., 2018). A similar phenomenon has been described for aldehyde dehydrogenase (ALDH) family member ALDH7A1 (Yang et al., 2019). GAPDH is also sensitive to oxidative stress and S-nitrosylation that causes its translocation to the nucleus where it can trigger apoptosis (Tristan et al., 2011). Thus along with effects on intermediary metabolism, consideration of metabolic enzyme subcellular distribution (and dynamics) and non-canonical protein-protein interactions will be required to truly understand the interface between intermediary metabolism machinery with neurodegeneration.

Oxidative stress is a cardinal feature of neurodegenerative disease in generally (Fransen et al., 2020; Lee et al., 2020) and ALS specifically (Barber et al., 2006) and direct targeting of reactive oxygen species (ROS) generation systems has been repeatedly shown to confer benefits in model systems (Greco et al., 2019; Sorce et al., 2017). The potent antioxidant Edaravone is one of two Food and Drug Administration approved medications for ALS (Abe et al., 2014). Endogenous antioxidants (i.e., the glutathione system, catalase and SOD1) all depend on the production of NADPH (Stanton, 2012). Since G6PDH is one of the major producers of NADPH it is perhaps counterintuitive that reducing its abundance or activity would be beneficial as we found. In addition to providing reducing power to antioxidant systems, G6PDH also has pro-oxidant activities – for example, both nitric oxide synthase and NADPH oxidase use NADPH to generate oxidants (nitric oxide and ROS, respectively) (Brennan et al., 2009; Leopold et al., 2003a; 2003b). Dysregulation of G6PDH has been show to drive reductive stress, protein aggregation and cardiomyotpathy in μB-Crystallin mutant mice (Rajasekaran et al., 2007). The opposing pro- and anti-oxidant activities of G6PDH may be linked to its dynamic subcellular localization (Stanton, 2012). While the metabolic effects of G6PDH in the setting of fALS mutant proteins is clearly not straightforward, we believe that deconstructing the role of G6PDH and pathological rewired metabolism in general will reveal novel targets for therapeutic intervention.

### Study Limitations

In this study we provided [1,2 ^13^C2]-D-glucose to neurons and monitored M2 lactate production to interrogate glycolysis and PPP. Several factors impact total and M2 lactate levels. For example, our measurements do not account for lactate generated from non-glycolytic sources (such as pyruvate recycling or oxidation of unlabeled amino acids) which would lower lactate APE. In addition, lactate formation and release to the medium occurs via concentration gradient-driven reactions, and the enormous extracellular compared with intracellular volume will ’pull’ lactate into the medium due to dilution, thereby increasing the apparent lactate production and glycolytic rates. In addition, M1 lactate production and calculated PPP rate would be underestimated when PPP intermediates are exported for nucleotide biosynthesis.

While neurons incubated with [1,2 ^13^C2]-D-glucose for 4 hours are very likely to have achieved isotopic steady state for glycolytic intermediates (Savaki et al., 1980; Veech et al., 1973), longer incubation times would be required to reach isotopic steady state of TCA cycle-derived amino acids (owing to their higher concentrations and longer half lives in brain in vivo). For comparison to M2 lactate, we report M2-labeled TCA cycle intermediates and TCA cycle-derived amino acids that are derived from the first turn of the TCA cycle. Subsequent turns of the cycle would produce additional isotopologues.

Potential isotopologue labeling differences among experimental groups can then arise from different TCA cycle rates and differences in metabolite retention within the cycle before label is incorporated into the amino acids. These possibilities were not examined in the present study.

## Acknowledgments

This work was supported by the National Public Health Service (NS05225 and NS087077), the Heather Koster Family Charitable Fund and the Les Turner ALS Foundation. We thank I. Nissim, Y. Daikhin, O. Horyn and I. Nissim for determining the ^13^C-enrichment and metabolite profiles in the Metabolomics Core Facility at the Children’s Hospital of Philadelphia; C. Link for sharing several strains of *C. elegans*, J. Rabinowitz for the gift of G6PDi-1, N. Chandel, R. Kibbey and I. Nissim for expert advice and members of the Kalb lab for fruitful discussions during the performance of these studies.

## Author Contributions

S.-P.R. performed all the *Saccharomyces cerevisiae, Caenorhabditis elegans* studies, analyzed data and created figures; J. M.-P. created all the mammalian neuron cultures and performed toxicity studies; M.G. performed *Caenorhabditis elegans* studies; V.M. and C.D. performed biochemical studies; G.D. analyzed and interpreted metabolomic data; J.M.-P. and R.G.K. performed all the metabolomic studies; R.G.K. supervised and integrated the project. All authors contributed to the writing and editing of the manuscript.

## Declaration of Interests

None

## STAR Methods

### Neuronal cultures

We began by designing an experimental platform in which heavy atom glucose could be used to monitor how neurons metabolize their major fuel. The following considerations guided our choice of this tissue culture experimental model. First, we needed sufficient material for analysis and only the embryonic cerebral cortex was a suitably abundant source of neurons to permit our detailed studies. Second, since neuronal metabolism is intimately linked to neuronal connectivity/network activity it was important to have a mixed neuronal population sufficiently mature that inter-neuronal communication could exist. Our system contrasts with patient-derived induced pluripotent stem cells (iPSC) that are differentiated into motor neurons, where such context-dependent factors are given less emphasis. Third, using viruses to express WT and mutant proteins allows for tight temporal control of toxic gene expression. This allows for a favorable level degree of biochemical synchrony that increases the sensitivity of our measurements. There are two main limitations to our system: 1) Since there are nominally no glial in our cultures, our model does not incorporate the metabolic cross talk between glial cells and neurons. This was a requirement for us because of our goal of biochemically interrogating neuronal metabolism specifically, and 2) WT and mutant proteins are heterologously expressed and thus potentially introducing non-physiological conditions. To mitigate these concerns, our studies are undertaken at an early time point when a robust metabolic phenotype was evident yet prior to any detectable neuron death(Mojsilovic-Petrovic et al., 2006).

Pure neuronal cultures were prepared using E17 rat cortex (meninges removed), and dissociated cells were grown on tissue culture plasticware coated with polylysine (50 nM) and laminin (3 ug/ml) in NeuroBasal, B27 and Glutamax. Fifty percent of media was refreshed 3x a week.

Spinal cord cultures were prepared as previously described (Mojsilovic-Petrovic et al., 2009) (Gupta et al., 2017; Zhai et al., 2015). In brief, astrocyte monolayers were generated from rat P1-3 rat cortex by dissociation and plated on German glass coverslips previously acid washed and coated with polylysine (50 nM) and laminin (3 ug/mL). When approximately ∼70% confluent, E15 rat spinal cord was dissociated and plated on top of the astrocytes in Minimal Essential Media with Earle’s salts (NEM) + 10% Horse serum (HS) with growth factors at 2 ug/ml ( CT-1, CNTF and GDNF). NEM + 10% HS is conditioned over astrocytes for one day prior to use. After 2 days, cyctosine arabinoside (AraC) was added (5 uM) to arrest the proliferation of astrocytes. Fifty percent of the growth media was replaced with fresh media three times a week and leading to progressive dilution of the AraC. For biochemical studies the same general approach was used except that astrocytes and neurons were grown on polylysine/laminin coated tissue coated plastic dishes.

### Metabolomic interrogations of primary neuron cultures

Metabolomic studies involved replacing standard media with an otherwise identical media in which all glucose was [1,2 ^13^C2]-D-glucose glucose and collecting media and cells 4 hours later for subsequent interrogations. We found that combining cells from two 60 millimeter (mm) dishes was compatible with our interrogations and a single experiment typically involved twenty-four 60 mm dishes.

### Biochemistry

Sample preparation for immune blotting was as previously described(Mojsilovic-Petrovic et al., 2009) (Gupta et al., 2017; Zhai et al., 2015). Briefly, cells were lysed in RIPA buffer (NaCl 150 mM, NP-40 1%, NaDeoxycholate 0.5%, SDS 0.1%, Tris pH 7.4 25 mM,), sonicated (15% duty cycle for 10s on ice), allowed to incubate for 30 minutes on a rotating drum at 4° C. before centrifugation at 16,000*g* in a microcentrifuge for 10 minutes. The supernatant was recovered, protein determined and western blot performed using denaturing SDS-PAGE gel.

To determine the efficacy of HSV engineered to express the miRNA targeting G6PDH, pure neuronal cultures were infected with recombinant virus and 48 hours later immunoblotted as described above.

### Immunocytochemistry

Pure neuronal cultures grown on glass coverslips were fixed with 4% paraformaldehyde for 10 – 20 minutes then washed extensively with PBS before incubating with primary antibodies overnight at room temperature (diluent was Dulbecco’s Modified Eagle Medium (DMEM) + 10% fetal bovine serum (FBS) + 1% triton X-100). The following day, coverslips were washed twice with PBS, incubated with species-specific Alexa fluor secondary antibodies (Invitrogen) for 4 – 6 hours as room temperature before washing in PBS twice, then water and mounting on slide with “Fluoro-Gel (with Tris buffer)”, Electron Microscopy Supplies. Hoechst 33342 (Thermo Scientific, diluted 1:1000 from manufacturers stock) was included with both primary and secondary antibody. Images were captures on a Leica DM14000 inverted confocal microscope with a 63x, 1.3 NA oil objective. Lasers and photomultipliers were set to non-saturating levels and single z-stack images are presented as typical examples of immunocytochemistry.

### Measurement of ^13^C-isotopogues

Measurement was performed in the Metabolomics Core facility of the Children Hospital of Philadelphia using either: (1) Liquid Chromatography- Mass Spectrometer (LC-MS) system, an Agilent Triple Quad 6410 MS combined with an Agilent LC 1260 Infinity: (2) Gas-Chromatograph-Mass Spectrometer (GC-MS) system, a Hewlett-Packard 5971 Mass Selective Detector (MSD), coupled with a 5890 HP-GC, and (3) GC-MS, an Agilent System 6890 GC-5973 MSD.

^13^C enrichment in the TCA cycle intermediates and/or in amino acids were performed as previously described (Li et al., 2014; 2008; Nissim et al., 2012). Briefly, cells were washed twice with PBS and then metabolites extracted with 4% perchloric acid (PCA).

Cell extracts were neutralized with KOH. The neutralized extracts were subjected to either AG-1 column (100-200 mesh, 0.5 x 2.5 cm, Bio-rad) for enriching the organic acids, or AG-50 (100-200 mesh, 0.5 x 2.5 cm, Bio-rad) for enriching amino acids. The collected samples were then converted to t-butyldimethylsilyl derivatives for GC-MS analysis. Enrichment in ^13^C glutamate isotopologues was monitored using ions at m/z 432, 433, 434, 435, 436 and 437 for M0, M+1, M+2, M+3, M+4 or M+5 (containing 1 to 5 ^13^C atoms above M0, the natural abundance), respectively. Isotopic enrichment in ^13^C GABA was monitored using ions at m/z 274, 275, 276, 277, and 278 for M0, M+1, M+2, M+3, or M+4 (containing 1 to 4 ^13^C atoms above M0, the natural abundance). Isotopic enrichment in ^13^C aspartate isotopologues was monitored using ions at m/z 418, 419, 420, 421 and 422 for M0, M+1, M+2, M+3 and M+4 (containing 1 to 4 ^13^C atoms above M0, the natural abundance), respectively. Isotopic enrichment in ^13^C lactate was monitored using ions at m/z 261, 262, 263 and 264 for M0, M1, M2 and M3 (containing 1 to 3 ^13^C atoms above natural abundance), respectively. Isotopic enrichment in ^13^C citrate isotopologues was assayed using ions at m/z 459, 460, 461, 462, 463, 464 and 465 for M0, M+1, M+2, M+3, M+4, M5 and M+6 (containing 1 to 6 ^13^C atoms above natural abundance), respectively. Isotopic enrichment in ^13^C malate isotopologues was determined using ions at m/z 419, 420, 421, 422 and 423 for M0, M+1, M+2, M+3 and M+4 (containing 1 to 4 ^13^C atoms above natural abundance), respectively. Isotopic enrichment in ^13^C fumarate isotopologues was monitored using ions at m/z 287, 288, 289, 290 and 291 for M0, M+1, M+2, M+3 and M+4 (containing 1 to 4 ^13^C atoms above natural abundance), respectively(Li et al., 2014; 2008; Nissim et al., 2012).

LC-MS system was used to determine isotopic enrichment in ^13^C-labeled ribose-5-P, and ^13^C-labeled acetylglucosamine. Protein was precipitated with PCA. The level of protein in 100ul was normalized to 1 ml for calculation of the relevant metabolite as amount/mg protein. Then, sample was derivatized by adding, 50 µl 0.3N NaOH, 50ul M 1-phenyl-3-methyl-5-pyrazolone (PMP), and placed into heater at 70°C for 30 min. After sample cooled, 50 µl of 0.3N HCL was added to the vial and mixed well. Then, 0.5 ml ultra-pure H2O and 2 ml ethyl acetate were added. Samples were vortexed for 1 min., centrifuged for 1 min. at 2500 RPM, the upper layer was removed and discarded, and the water phase was evaporated completely. The dry sample was reconstituted with 100 µl 0.1% formate in water (Solution A), vortexed, transferred into injection vials, and 3 µl was injected into LC-MS. Chromatographic conditions were as follows: Separation was performed on Agilent Poroshell 120 EC-C18 column. The mobile phase consists of Solution A and Solution B (0.1% formate in acetonitrile mixed with 0.005% trifluroacetate). LC flow was directed into waste for the first 1.5 minute, and then, diverted into MS for next 6 min. and back to waste at 7.5 min to flush the column, and stop time of 10 min. MS condition were: Capillary voltage, 4000V, nebulizer 25 p.s.i, and dry gas temperature at 350°C. Fragmentor and Collision energy voltage were established for each compound by MassHunter Optimizer software. Metabolites were monitored using multiple-reaction monitoring (MRM) in positive mode. For ribose-5-P, at retention time (RT) 3.7 min., we monitored M0, M+1 and M+2 using 561-175,562-175 and 563-175 MRM, respectively. For acetylglucosamine, at RT 4.9 min. we monitored M0, M+1 and M+2 using 552-175,553-175 and 554-175 MRM, respectively. For glucosamine, we monitored M0, M+1 and M+2 using 510-175, 511-175 and 512-175 MRM, respectively. Glucosamine, if present in the sample, was below the detection limit, and, therefore, not reported in the Results Section.

### Measurement of metabolite levels

The levels of media glucose and its net uptake during the incubation was determined as described (Li et al., 2014; Nissim et al., 2012; 2014) and the results were expressed as μmol/mg cellular protein. The levels of ATP, ADP and AMP were measured as in (Li et al., 2014; Nissim et al., 2012; 2014). The levels of NADP+ and NADPH were determined according to the manufacture’s protocol using the BioVision Colorimetric Kit (Catalog#K347). The levels of NAD^+^ and NADH were determined according to the manufacture’s protocol using the BioVision Colorimetric Kit (Catalog#K337). The concentrations of amino acids were determined with an Agilent 1260 Infinity HPLC system, utilizing pre-column derivatization with O-phthalaldehyde (Nissim et al., 2012) (Nissim et al., 2014). Lactate levels were determined enzymatically as in (Li et al., 2014; Nissim et al., 2012; 2014). Concentrations of representative of the TCA cycle intermediates (fumarate, citrate and succinate) in cell extracts were determined by isotope dilution approach using GC-MS system as described (Weinberg et al., 2000).

Briefly, an aliquot of the sample (100 μl) was spiked with a mixture of ^13^C-labeled organic acids of known concentrations. Then, GC-MS measurement of ^13^C isotopic abundance in each sample was performed (GC-MS parameters and measurements as described above, and the concentrations in the sample were calculated as described (Weinberg et al., 2000).

Concentrations of GSH and GSSG were determined by isotope dilution approach using N-ethylmaleimide (NEM) derivatization and LC-MS system. Method details are as in (Moore et al., 2013). Briefly, cell pellets were spiked with a known concentration of [^13^C2,^15^N]-GSH (Sigma) and GSH-ethylester (GSH-EE, Sigma) which were used as an internal standard. Then, sample was subjected to three freeze/thaw cycles in 50ul of 240mM NEM in15% Methanol, vortexed vigorously, and incubated for 1 hour at room temperature. Incubation was followed by addition of 50 µl of precipitating solution that contained 240mM NEM, 10% sulfosalicylic acid, 2mM EDTA and 15% methanol.

Samples were vortexed vigorously, centrifuged for 5 min. at 14000 RPM, the supernatant was transferred to LC-MS injection vials, and samples injected to LC-MS; the conditions of LC and MS were similar to above. The RT of GSH 3.9 min. and GSSG 3.6 min., we used 436-307 MRM to monitor ^13^C-^15^N-GSH, 461-332 MRM to monitor GSH-EE, 433- 304 MRM to monitor GSH and 613-355 MRM to monitor GSSG. Calculation of the concentrations were determined as in (Weinberg et al., 2000). All measurements described above were validated by standard curves derived from pure metabolites.

Cellular protein levels were determined with Coomassie blue (Thermo Fisher Scientific) (Nissim et al., 2012), (Nissim et al., 2014).

### Calculations and Statistical Analysis

^13^C-enrichment in ^13^C-labeled mass isotopomers is expressed by Atom Percent Excess (APE), which is the fraction (%) of analyte containing ^13^C atoms above natural abundance. The amount of ^13^C-labeled mass isotopomer was calculated by the product of (APE/100) times concentration (nmoles **x** mg^-1^ cellular protein) and is expressed as nmoles ^13^C metabolite per mg cellular protein. The flux of the relative pentose phosphate (RPP) was quantified by the ratio of M+1 to M+2 in ^13^C-labeled lactate and overall flux through pentose phosphate pathway (PPP) was calculated as described (Lee et al., 1998). CMRglc[(M+1)lactate/(M+1 + M+2)lactate], where M+1 and M+2 denote the ^13^C contents of lactate: (M+1)lactate represents lactate derived from the PPP due to decarboxylation of ^13^C-labeled carbon one of [1,2-^13^C2]glucose-6-phosphate, and (M+2)lactate represents lactate derived from the glycolytic pathway.

Overall rates of glucose utilization (CMRglc) were calculated by dividing net glucose uptake by the 4h incubation interval. Overall rate of lactate release was calculated as net increase in medium lactate concentration divided by the 4h experimental interval, and the rate was converted to glucose release equivalents by dividing by 2. The PPP was converted to the same units as CMRglc, and all rates are expressed as micro-mol glucose or its equivalent/mg/h so they can be compared directly. For data summaries, rates were expressed either % of CMRglc or as % of the respective rate in LacZ constructs (Figure 2 M. – P.). For the 100% values for LacZ cells, the error bars for the rate calculations are coefficients of variation cv= 100*SD/mean. For all other values, the SDs were calculated as using the following equation that accounts for error propagation and expressed as percentages.

### Yeast strains

The rationale for selecting yeast strains for these studies were guided by three considerations. First, there are often multiple Isoforms (encoded by separate genes) of particular glycolytic enzymes that differ in intrinsic activity, localization and responsiveness to environmental conditions (Jang et al., 2016; Li et al., 2020b; van Geersdaele et al., 2013; Zala et al., 2013; Zhang et al., 2014). Second, in addition to energy production *per se*, metabolic intermediates can play crucial roles in a variety of cell biological processes (i.e., senolysis (Johmura et al., 2021), TOR modulation (Chin et al., 2014) and chromatin regulation (Gut and Verdin, 2013)). Third metabolic enzymes can participate in protein-protein interactions that modify cellular processes independent of their enzymatic activity (Li et al., 2014; Yang et al., 2018). For these reasons, it may be difficult to predict the effect of ablation of a specific gene on both substrate flux and molecular cell biology in general. Thus it was important to adopt an approach that was not constrained by prior understanding of the function of specific metabolism enzyme.

All yeast strains in this study are taken from the publicly available yeast deletion library (distributed by Horizon Discovery) (Supplemental Table 3). In case of deletion mutants of essential genes, we used BY4743 (MATa/α; *his3Δ1/his3Δ1; leu2Δ0/leu2Δ0; met15Δ0/MET15; LYS2/lys2Δ0; ura3Δ0/ura3Δ0*) based strains whereas we used BY4741 (MATa *his3Δ1 leu2Δ0 met15Δ0 ura3Δ0*) based deletion strains for non-essential genes (Brachmann et al., 1998). All yeast strains were checked by colony PCR for corresponding deletions before use.

### Yeast Plasmids

All vectors used in this study were generous gifts from James Shorter. cDNAs of ATXN3 Q25 and ATXN3 Q71 were generous gifts of Erich E. Wancker. In the plasmids the expression of proteins of interest (TDP-43, TDP-43 Q331K, TAF15, EWSR1, Fus, Fus R521C, Fus P525L, α-Syn, α-Syn E46K α-Syn A53T, ATXN3 Q25, ATXN3 Q71) is under the control of a galactose inducible promoter. Vectors (pRS423Gal, pRS413Gal) were originally modified from redesigned pRS vectors (Chee and Haase, 2012).

### Yeast transformation

Yeast strains were transformed using the lithium-acetate method modified from Ito *et al*. (Ito et al., 1983). Briefly, 1 ml of yeast overnight cultures in YPD define was used for inoculating 28 ml of fresh YPD in a 250 ml Erlenmeyer flask. Cells were harvested for 5 min at 1,000 g after about 4 h after reaching an OD600 of 0.6-0.8. Cells were washed in 10 ml of 1x TE buffer, pH8. Cells were made competent by suspending them in 1 ml of TE based 0.1 M lithium acetate solution and incubating them for 10 min at room temperature. On wet ice, 0.5-2 µg plasmid DNA was mixed with 5 µl of carrier DNA (Clontech Order No.: 630440) and 12 µl of competent yeast cells. 60 µl of PEG-solution (0.1 M lithium acetate and 60% PEG 8,000 in 1x TE buffer) were added and mixed gently. The transformation reaction was incubated at 30°C for 30 min before adding 8 µl of DMSO. After that, cells were heat shocked for 7 min at 42°C. Cells were applied to selective agar plates (SD -His) and checked for transgenic colonies after 4-7 days at 30°C. Finally, single colonies were selected and verified by colony PCR.

### Yeast disease gene toxicity modifier assay

The assay was performed as described earlier (Armakola et al., 2012) (Couthouis et al., 2011) with modifications. Briefly, yeast cells were grown for 3 d at 30°C in liquid medium with glucose (SD –His). After that, cells were grown for 17 h at 30°C in liquid medium containing raffinose (SRaf –His). Cultures were harvested and re-suspended in YNB without amino acids adjusted to an OD600 of 100. Cells of different samples were spotted in identical serial dilutions on plates containing either galactose (SGal –His) for disease protein expression or glucose (SD –His) as a non-expression control. Cells on plates were grown for 2 d before photos were taken for data analysis. Spots were evaluated and ratios were calculated for number of spots with growth with disease protein expression in relation to number of dilution steps with growth without disease protein expression. After that, growth ratios with and without a deletion for a metabolic gene were compared to each other (Figure 4, A.). Experiments were done in 2-4 biological replicates with 3 technical replicates each.

### C. elegans strains

CL6303 *(eri-1(mg366) IV; uls60[P-unc-119::YFP +P-unc-119::sid-1]; dvIs62(snb- 1::TDP-43)X*, CL6186 - eri-1(mg366) IV), and CL6264 *(uls60[P-unc-119::YFP +P-unc-119::sid-1]*) were a generous gift of Christopher D. Link. Strain RK179 (*eri-1(mg366) IV; uls60[P-unc-119::YFP +P-unc-119::sid-1]*) was constructed and genotyped using CL6186 and CL6264 as parental strains.

### Worm cultivation

*C. elegans* were cultivated at 20 °C on nematode growth medium (NGM) agar surface, unless otherwise stated, using standard techniques (Brenner, 1974). NGM plates were seeded with 50 µl of the *E. coli* strain OP50 as the food source. *E. coli* cells in general were grown for 1-2 d and formed a lawn before placing worms on plates.

### Knock-down of genes of interest in worms

For feeding RNAi experiments NGM plates were seeded with *E. coli* (HT115 (F-, mcrA, mcrB, IN(rrnD-rrnE)1, rnc14::Tn10(DE3 lysogen: lacUV5 promoter -T7 polymerase)) expressing empty vector (L4440) or different RNAi-encoding plasmids (Supplemental Table 4). We tested the worm strain CL6303 and RK179 strains, enhanced for neuronal import of dsRNA from the bacterial source and RNAi-dependent silencing of the target gene, to achieve knockdown of the genes of interest specifically in neurons. L4-staged worms were placed on either empty vector or gene of interest RNAi plates (NGM with 25 µg/ml Carbenicillin and 1 mM isopropyl β-D-1-thiogalactopyranoside (IPTG)) for 2–4 generations to ensure successful knockdown of the gene of interest. Subsequently, L4 staged and young adult worms were used in the worm fitness assay.

### Worm swimming assay

Worms were grown for 2-4 generations either on HT115 carrying either L4440 empty vector or L4440 expressing the RNAi for a gene of interest. After that, 9-15 L4 stage and young adult worms were transferred onto a RNAi plate without *E. coli* cells and suspended in a drop M9 buffer. After that, the animals were tested immediately in a worm swimming assay as described earlier using the ImageJ plugin wrMTrck (Nussbaum-Krammer et al., 2015). Briefly, 1 minute movies were taken using bright- field microscopy with a high-resolution CCD camera. We obtained 6-14 movies of each knock-down tested. Using the wrMTrck plugin readout we determined the average speed, fold <u>c</u>hange (FC), the average bending rate FC, average path length, and the average distance traveled by an animal per minute. FC’s were calculated as values when a metabolism gene was knocked down in relation to the empty vector control.

### miR RNA design for the knock-down of glycolysis genes in rat primary neurons

All miR RNA’s in this paper were designed using the BLOCK iT^TM^ RNA designer (https://rnaidesigner.thermofisher.com/rnaiexpress/). Briefly, miR RNA was selected as target design option, the rat (*Rattus norvegicus*) open reading frame (ORF) was used for miR RNA design, minimum G/C content of 35% and maximum of 55% were selected. The three top ranking sequences selected by the design tool that do not overlap in their position in the ORF were selected. The tool designs a top strand as well as a bottom strand. For cloning the miR RNA constructs into the p1006 (+) empty vector using the *Fse*I and *Bsp*EI restriction sites, we added 5’ and 3’ flanking sequences to the top strands (5’: CCTGGAGGCTTGCTGAAGGCTGTA; 3’: CAGGACACAAGGCCTGTTACTAGCACTCACATGGAACAAATGGCCT), as well as 5’ and 3’ flanking sequences to the bottom strands (5’: CCGGAGGCCATTTGTTCCATGTGAGTGCTAGTAACAGGCCTTGTGTCCTG; 3’: AGCATACAGCCTTCAGCAAGCCTCCAGGCCGG). The final top and bottom oligonucleotides were purchased from IDT and dissolved in nuclease free water at 200 µM. 4 nmol of top and bottom oligonucleotides each were mixed in annealing buffer (10 mM Tris-HCl, pH8.0, 1 mM EDTA, 100 mM NaCl) and heated to 95°C for 4 min, followed by cooling for 10 min at room temperature. The annealed double-stranded oligonucleotide was diluted to 10 nM in annealing buffer. In parallel the HSV p1006 (+) vector was digested using *Fse*I and *Bsp*EI (both NEB) and the linearized vector was gel purified. The *Fse*I/*Bsp*EI linearized HSV p1006 (+) (2.5 ng/µl) and the double-stranded miR RNA construct (4 nM) were ligated using the T4 fragment as recommended by the manufacturer for 30 min at room temperature and transformed into chemically competent *E. coli* cells (TOP10; Thermo). Finally, single colonies were picked, purified (Qiagen), sequence verified (NU sequencing facility) and tested for knock-down efficiency of heterologous expressed Flag-tagged rat G6pd in HEK293 cells in Lipofectamine 2000 (Thermo) transfections as recommended by the manufacturer.

Constructs showing efficient target construct knock-down were packed in HSV viral vectors at the Massachusetts General Hospital Gene Delivery Technology Core on a fee for service basis.

### Statistical analysis

Data were analyzed using Prism (GraphPad Software, La Jolla, CA). Significant differences between two groups were determined using paired Student’s *t*-test (two-tailed). Significant differences within groups greater than two were determined using one-way ANOVA followed by Tukey’s test for multiple comparisons. Data are presented as mean ± SD. For all tests, the significance threshold was set to p < 0.05

### Inclusion and diversity Statement

We worked to ensure sex balance in the selection of subjects and experimental samples throughout this project. None of the authors of this manuscript self identify as an underrepresented ethnic minority, a member of the LGBTQ+ community, living with a disability or received support from a program designed to increase minority representation in science. We actively worked to promote gender balance in our reference list.

### Data and Image processing

Data processing was kept to a minimum and conform to the guidelines provided to authors on the Cell Metabolism website.

### Studies involving human and animals

No human samples were used in this study and the use of vertebrate animals was covered by our approved Animal Care and Use Committee Protocol.

### Resource availability

All yeast and worm strains used in this study are freely available to qualified investigators by simple request. In addition, plasmids used for the creation of viruses (amplicons) are similarly freely available to qualified investigators by simple request.

### Data and Code Availability

All primary data collected in the course of these experiments is freely available to qualified investigators by simple request.

## KEY RESOURCES TABLE

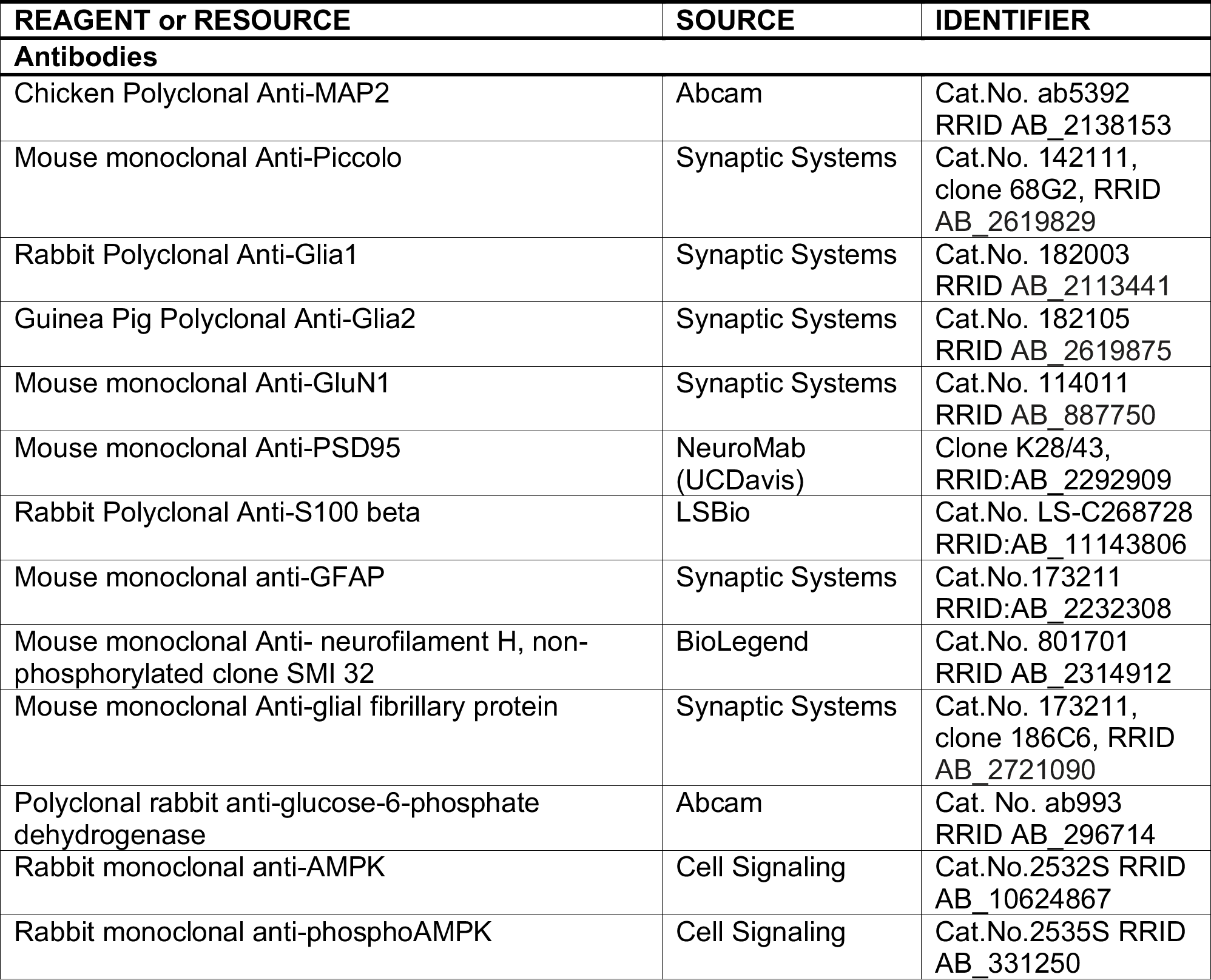
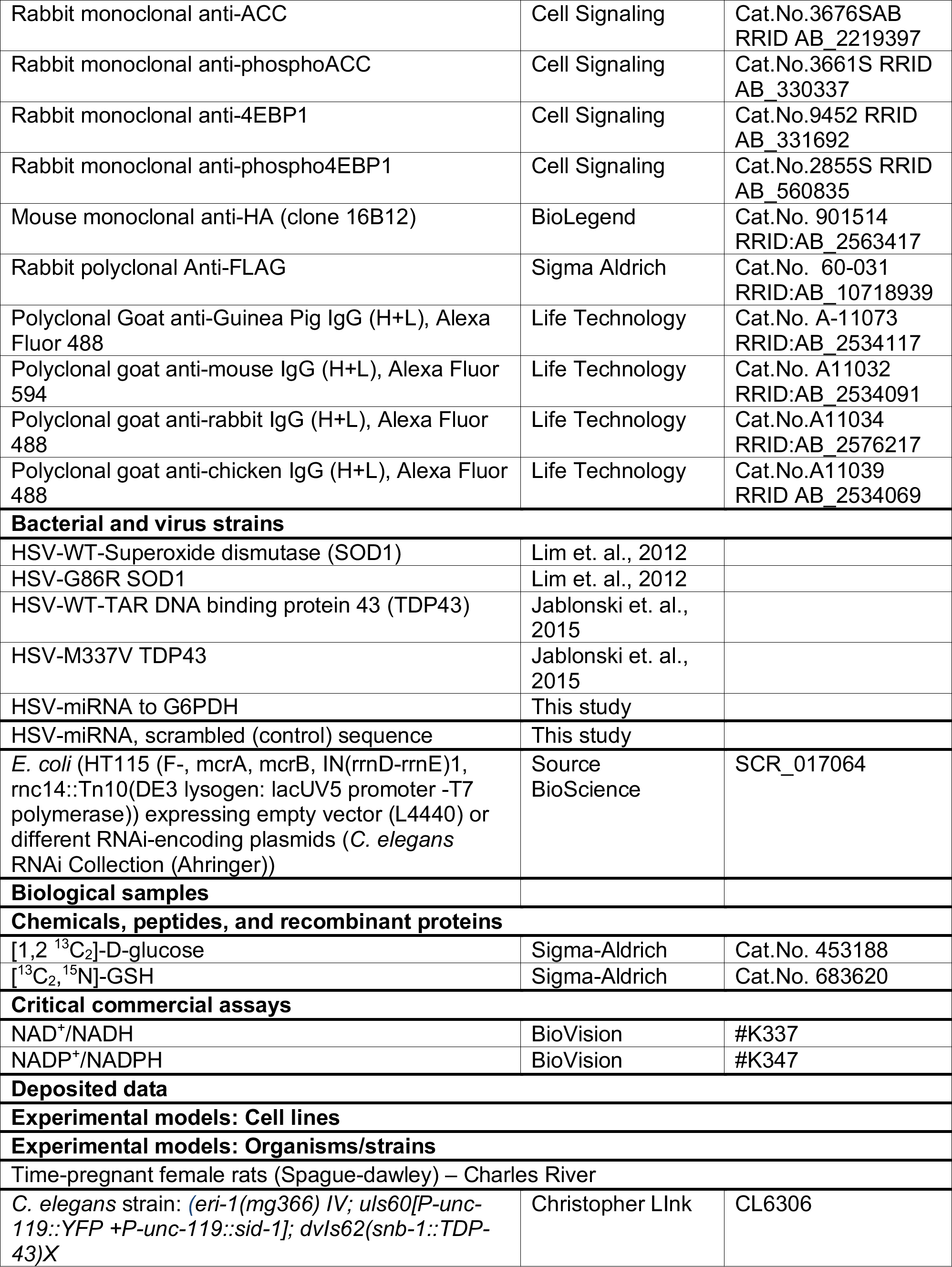
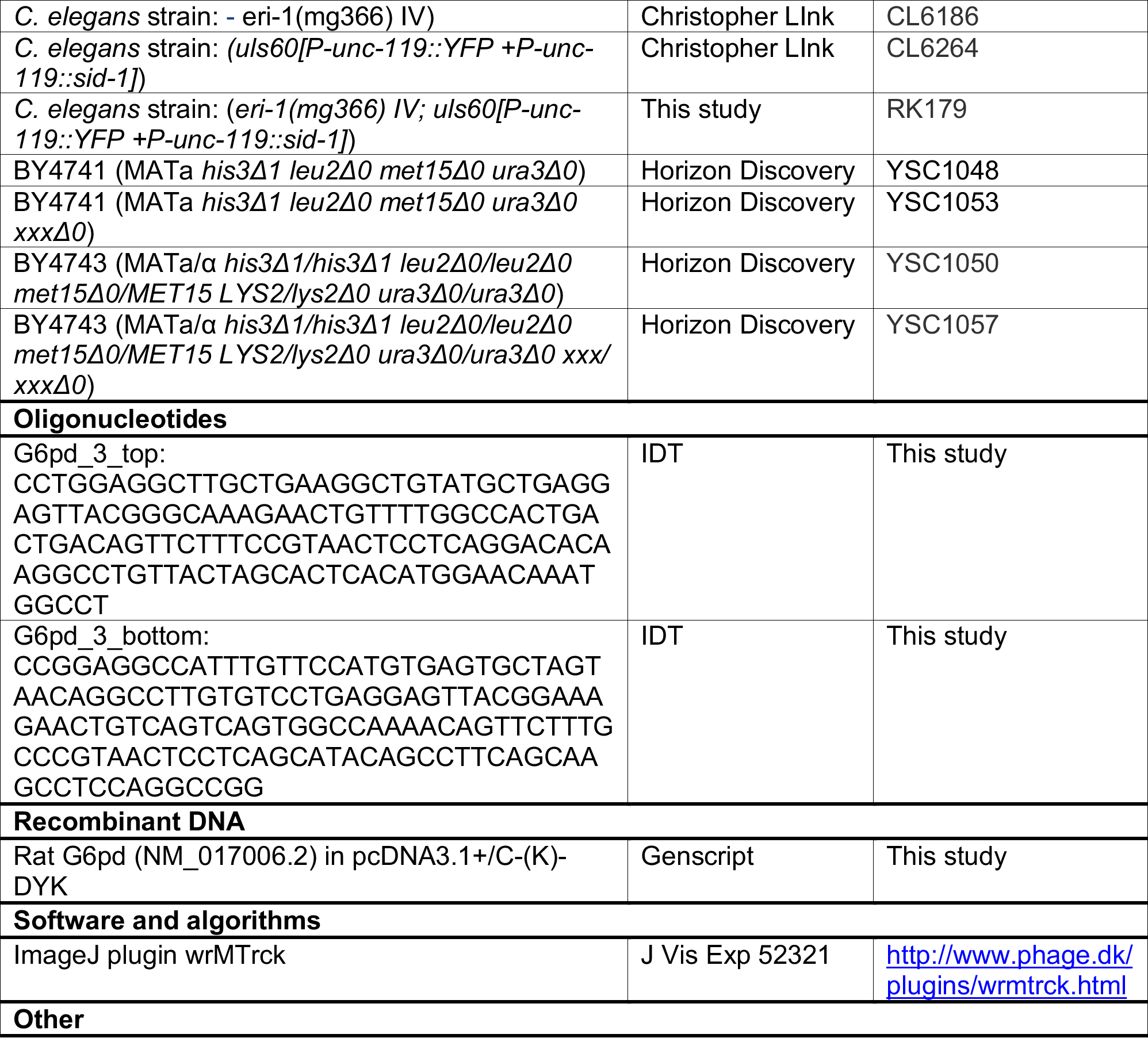

## Supplemental Information titles and legends

**Supplemental Figure 1.**
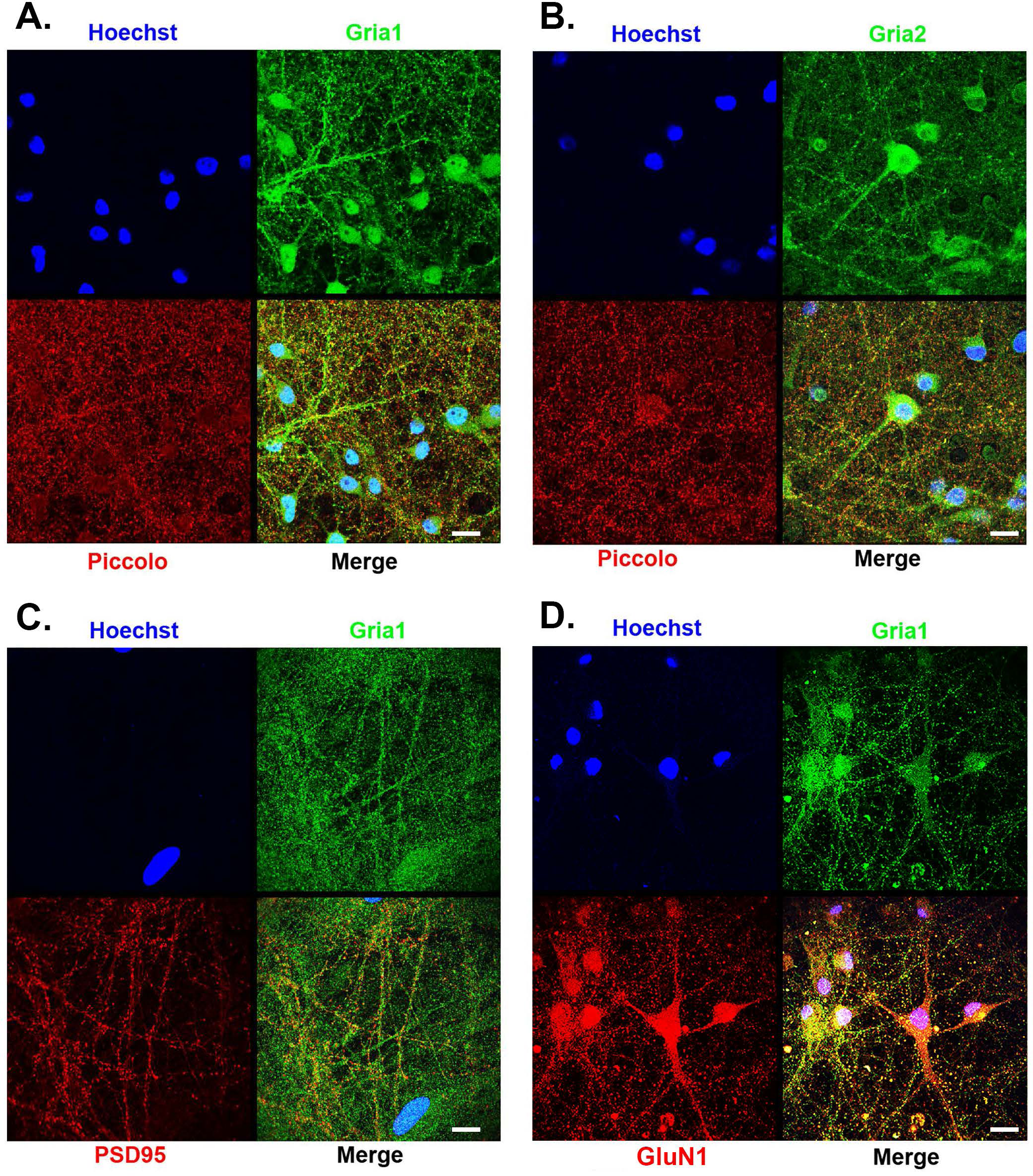
Immunocytochemically-defined pre- and post-synaptic elements indicate that there is synaptic connectivity in cortical neuron cultures. Panel A. DIV 14 cortical neuron cultures underwent immunocytochemical staining for Hoechst staining of nuclei, the Gria1 subunit of AMPA receptors (green) and the presynaptic protein, Piccolo (red). Merge image of a single confocal slice reveals pre- and post-synaptic apposition of puncta reflecting the existence of AMPA’ergic synapses. Calibration bar, 30 microns. Panel B. DIV 14 cortical neuron cultures underwent immunocytochemical staining for Hoechst staining of nuclei, the Gria2 subunit of AMPA receptors (green) and the presynaptic protein, Piccolo (red). Merge image of a single confocal slice reveals pre- and post-synaptic apposition of puncta reflecting the existence of AMPA’ergic synapses. Calibration bar, 30 microns. Panel C. DIV 14 cortical neuron cultures underwent immunocytochemical staining for Hoechst staining of nuclei, Gria1 subunit of AMPA receptors (green) and the postsynaptic density protein, PSD95 (red). Merge image of a single confocal slice reveals Gria1 puncta adjacent to PSD95 reflecting the existence of AMPA’ergic synapses. Calibration bar, 30 microns. Panel D. DIV 14 cortical neuron cultures underwent immunocytochemical staining for Hoechst staining of nuclei, the Gria1 subunit of AMPA receptors (green) and GluN1 subunit of NMDA receptors (red). Merge image of a single confocal slice reveals Gria1 puncta co-localizing with GluN1 puncta reflecting the existence of synapses containing both AMPA and NMDA receptors. Calibration bar, 30 microns.

**Supplemental Figure 2.**
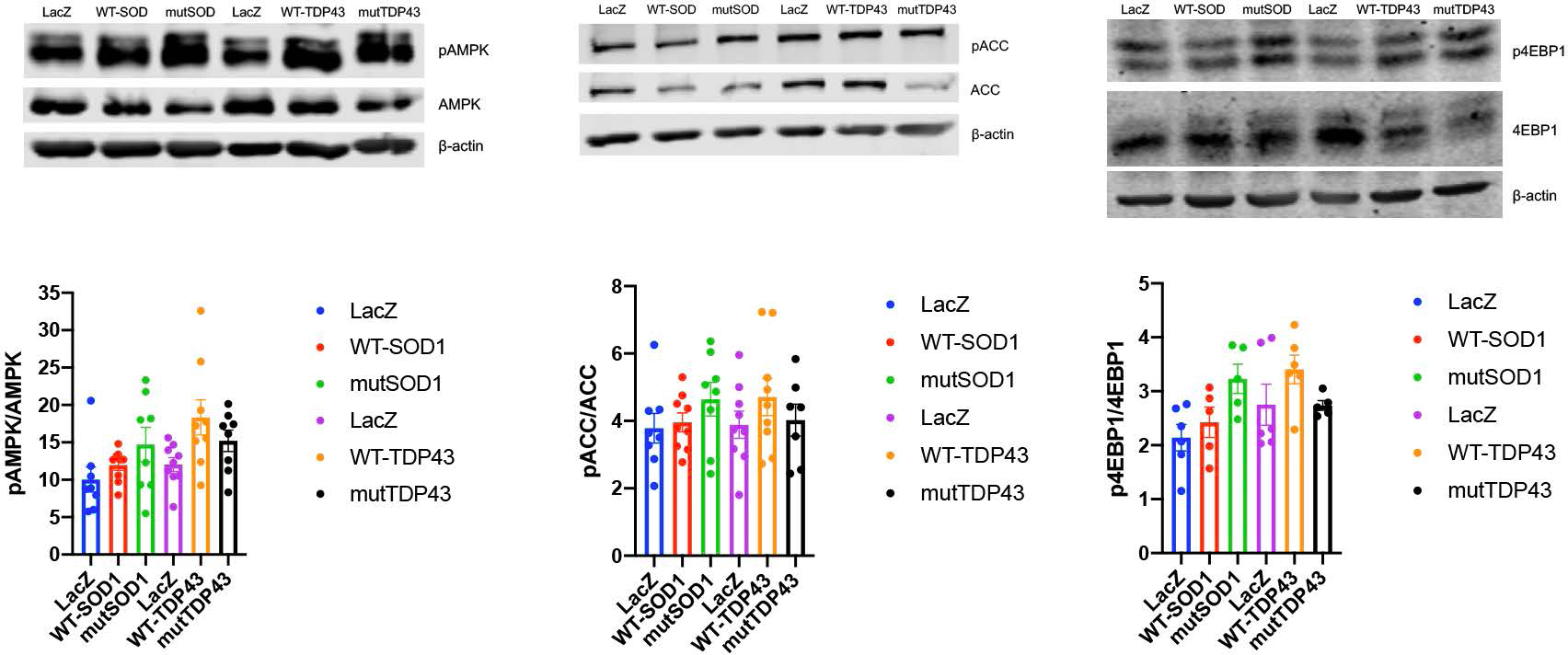
AMPK is not activated in neurons expressing fALS genes. DIV14 pure neuronal cultures were infected with HSV engineered to express LacZ, WT or mutant form of SOD1 or TDP43 and two days later Western blots were performed on lysates. AMPK activation is monitored by blotting for phosphorylation of threonine 172 (e.g., pAMPK) and the state of phosphorylation of downstream targets ACC (phosphorylated serine 79, pACC) and 4EBP1 (phosphorylated threonine 37/46, p4EBP1). Representative blots are displayed and quantification of pAMPK/AMPK, pACC/ACC and p4EBP1/4EBP1 presented below. There was no statistically significant difference between the ratio of phosphorylated to non-phosphorylated species for any of these measures of AMPK activation by neurons expressing WT or mutated fALS proteins versus LacZ. These results were derived from 6 – 9 independent pure neuron cultures.

**Supplemental Figure 3.**
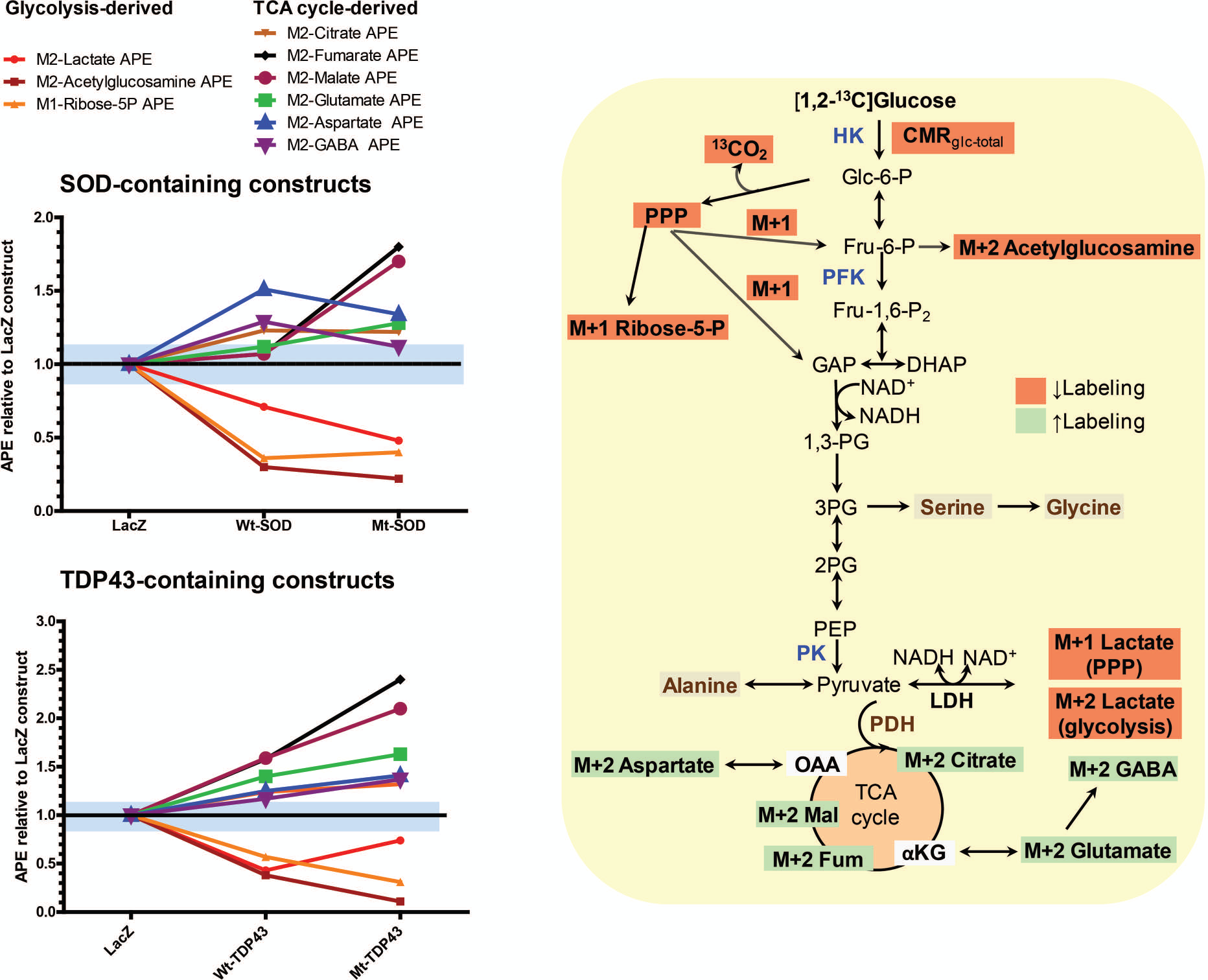
Experimental ALS causes upregulation of flux of [1,2-^13^C2]glucose into oxidative pathways and downregulation of glycolysis and its branch pathways. Mean values for the atom percent excess (APE, i.e., the fraction of the analyte containing ^13^C atoms above natural abundance; see Methods ) for each metabolite were normalized to the respective values for the LacZ constructs and plotted for each experimental group. The individual and mean values for each experimental group and statistical differences among groups are presented in Figures 2 and 3 of the main text. The blue shaded area represents ± mean 95% confidence interval (CI) that was calculated from the 95% CIs for all compounds in the LacZ constructs. Metabolites produced by (i.e., lactate) or derived from branch pathways (i.e., ribose-5-phosphate (P) and acetylglucosamine) of the glycolytic pathway (pathways are illustrated in the right panel) have reduced enrichment from [1,2-^13^C2]glucose in the wildtype (Wt) and mutant (Mt) Cu^++^/Zn^++^ superoxide dismutase (SOD1) (top left panel) and TAR DNA binding protein of 43 kDa molecular weight (TDP43) (bottom left panel) constructs compared with LacZ constructs. In contrast, metabolites that are components of the tricarboxylic acid (TCA) cycle (fumarate (Fum), malate (Mal), and citrate (Cit)) and amino acids derived from the TCA cycle (aspartate, glutamate, and gamma-aminobutyric acid (GABA)) have increased enrichment compared with the LacZ construct. Other abbreviations: M+1 or M+2, mass of the compound containing 1 or 2 ^13^C atoms derived from [1,2-^13^C2]glucose, respectively; HK, hexokinase; Glc-6-P, glucose-6-P; PPP, pentose phosphate pathway; Fru-6-P, fructose-6-P; PFK, phosphofructo-1-kinase; Fru- 1,6-P2, fructose-1,6-bisphosphate; GAP, glyceraldehyde-3-P; DHAP, dihydroxyacetone-P; 1,3-PG, 1,3-diphosphoglycerate; 3PG, 3-phosphoglycerate; PEP, phosphoenolpyruvate; PK, pyruvate kinase; LDH, lactate dehydrogenase; PDH, pyruvate dehydrogenase; OAA, oxaloacetate; μ- KG, μ-ketoglutarate.

**Supplemental Figure 4.**
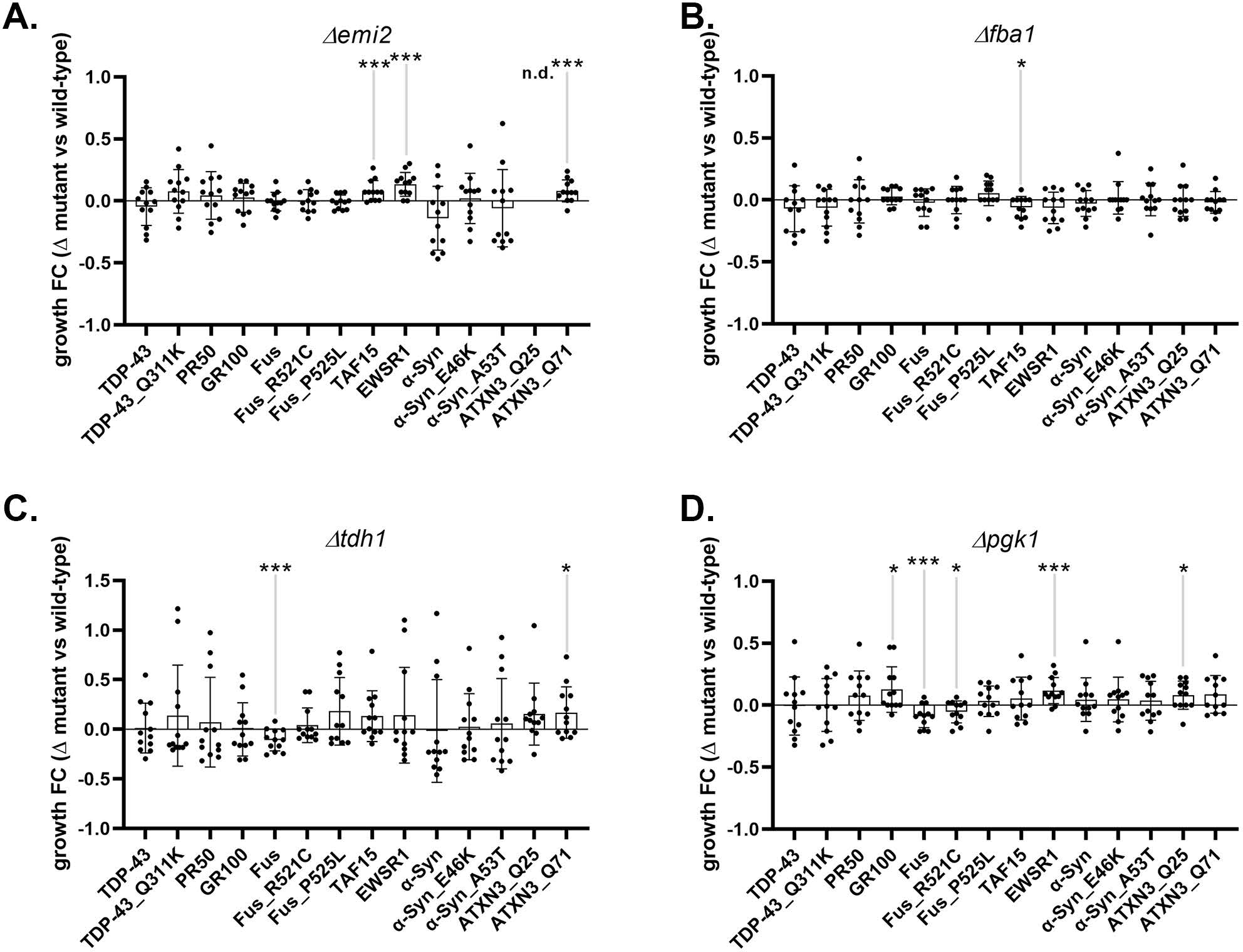
Deletion of individual glycolytic enzymes reveal modifiers of protein toxicity of neurodegenerative disease-associated proteins in yeast. Panel A. Growth fitness effects of ablation of *emi2* on a variety of proteotoxic insults relevant to ALS, PD and SCA; n=12. Growth fitness benefits were seen in yeast expressing TAF15, EWSR1 and ATXN3 Q71. Panel B. Growth fitness effects of ablation of *fba1* on a variety of proteotoxic insults relevant to ALS, PD and SCA; n=12. Growth fitness benefits were seen in yeast expressing TAF15. Panel C. Growth fitness effects of ablation of *tdh1* on a variety of proteotoxic insults relevant to ALS, PD and SCA; n=12. Growth fitness benefits were seen in yeast expressing ATXN3 Q71. Growth fitness detriment was seen in yeast expressing FUS. Panel D. Growth fitness effects of ablation of *pgk1* on a variety of proteotoxic insults relevant to ALS, PD and SCA; n=12. Growth fitness benefits were seen in yeast expressing GR100, EWSR1, ATXN3 Q25. Growth fitness detriment was seen in yeast expressing FUS and FUS R521C (*p<0.05, ***p<0.005).

**Supplemental Figure 5.**
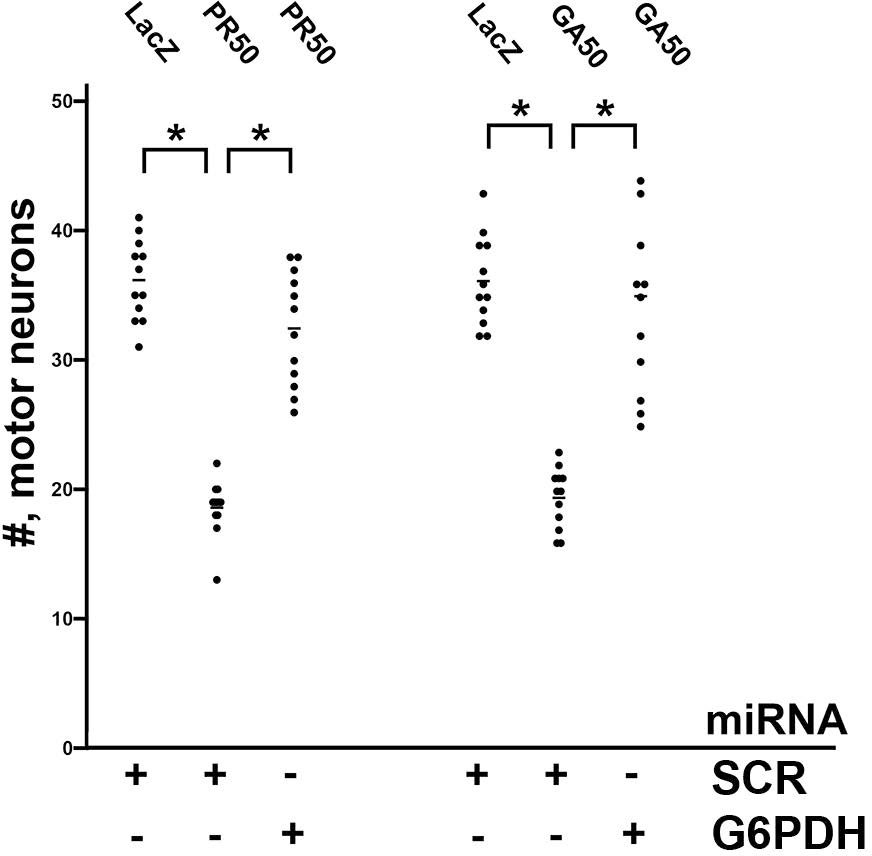
Reduction of G6PDH abundance protects against the motor neuron toxicity of PR50 and GA50. Panel A. A three way comparison of number of motor neurons expressing LacZ + scr, PR50 + scr miRNA and PR50 + G6PDH miRNA. Statistically significant group differences were seen by ANOVA and post hoc test reveal that PR50 + scr miRNA is significant different from LacZ +scr and from PR50 + G6PDH miRNA. Results from ≥4 experiments, *, p < 0.05. A three way comparison of number of motor neurons expressing LacZ +scr, GA50 + scr miRNA and GA50 + G6PDH miRNA. Statistically significant group differences were seen by ANOVA and post hoc test reveal that GA50 + scr miRNA is significant different from LacZ +scr and from GA50 + G6PDH miRNA. Results from ≥4 experiments, *, p < 0.05.

**Supplemental Figure 6.**
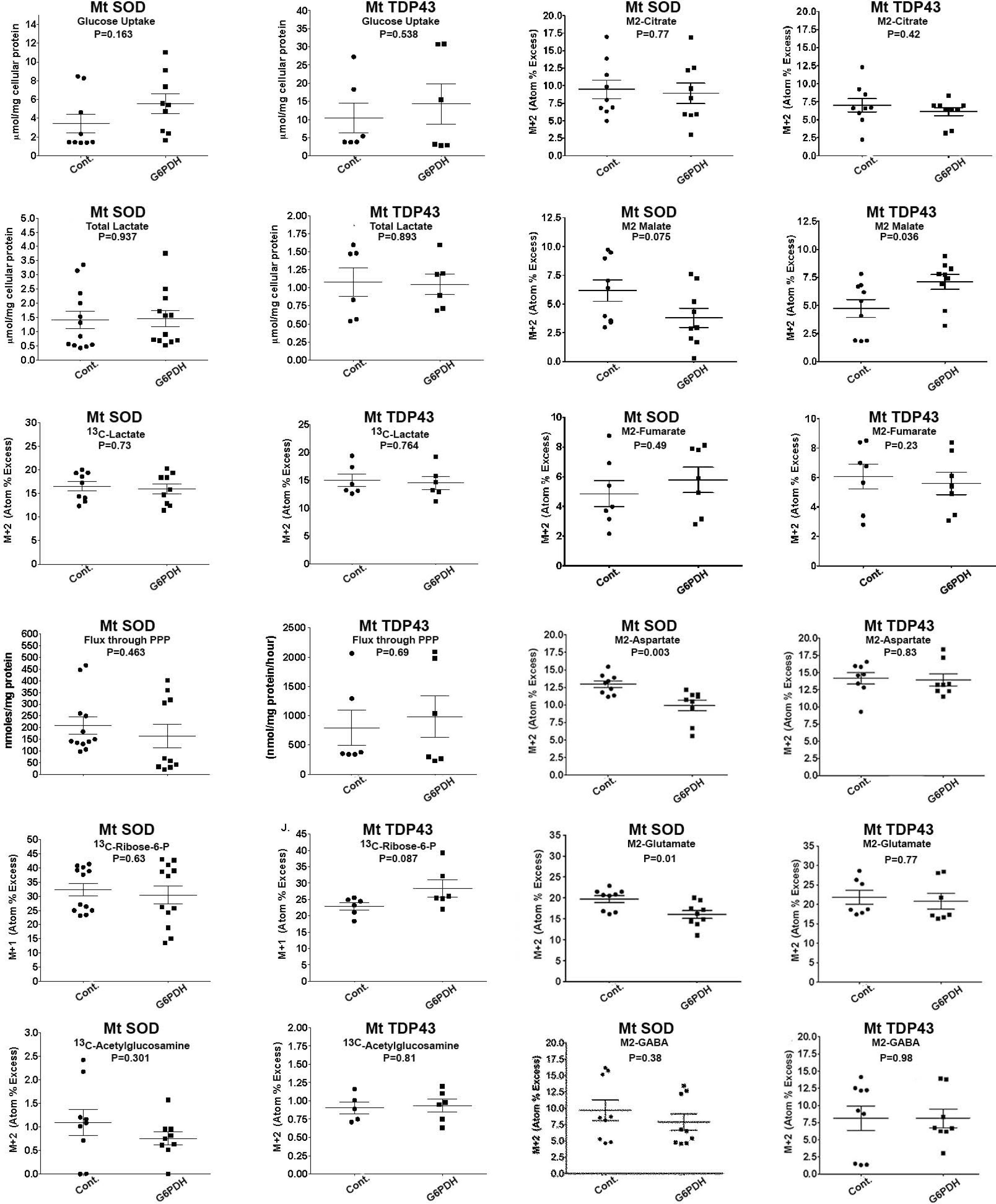
Metabolic interrogation of glycolysis, theTCA cycle and related pathways in neurons expressing fALS genes with or without knockdown of G6PDH. Scattergrams of data from neurons expressing Mt-SOD1 or Mt TDP43 with miRNA mediated knockdown of G6PDH or expression of a control miRNA (”Cont.”). P values from Student’s t-test are presented in every panel. The only statistically significant differences were: 1) higher levels of M2-malate in Mt TDP43 expressing neurons with G6PDH knockdown versus control, 2) lower levels of M2-asparate in Mt SOD1 expressing neurons with G6PDH knockdown versus controls and 3) lower levels of M2- glutamate in Mt SOD1 expressing neurons with G6PDH knockdown versus controls (compare to Figures 2 and 3 in main text).

**Table 1.** Total intracellular amino acid levels from neuronal cultures engineered to express LacZ, WT or mutant versions of SOD1 or TDP43. Mean, standard error and ANOVA with *post hoc* analysis are presented.

| <u>ANOVA Analysis</u> |  |  |  |
| --- | --- | --- | --- |
|  | <u>Wt-TDP43vs. LacZ</u> | <u>Mt-TDP43 vs. LacZ</u> | <u>WT-TDP43 vs. MT-TDP43</u> |
| ASP | NS | NS | NS |
| GLU | NS | NS | NS |
| SER | NS | NS | NS |
| GLN | 0.001 | NS | NS |
| HIS | NS | NS | NS |
| GLY | NS | NS | NS |
| THR | NS | NS | NS |
| ARG | NS | NS | NS |
| TAU | 0.001 | NS | NS |
| ALA | NS | NS | NS |
| GABA | NS | NS | NS |
| TYR | NS | NS | NS |
| METH | NS | NS | NS |
| VAL | NS | NS | NS |
| PHE | NS | NS | NS |
| IL | NS | NS | NS |
| LEU | NS | NS | NS |
| ORN | NS | NS | NS |
| LYS | NS | NS | NS |

**Table 2.**
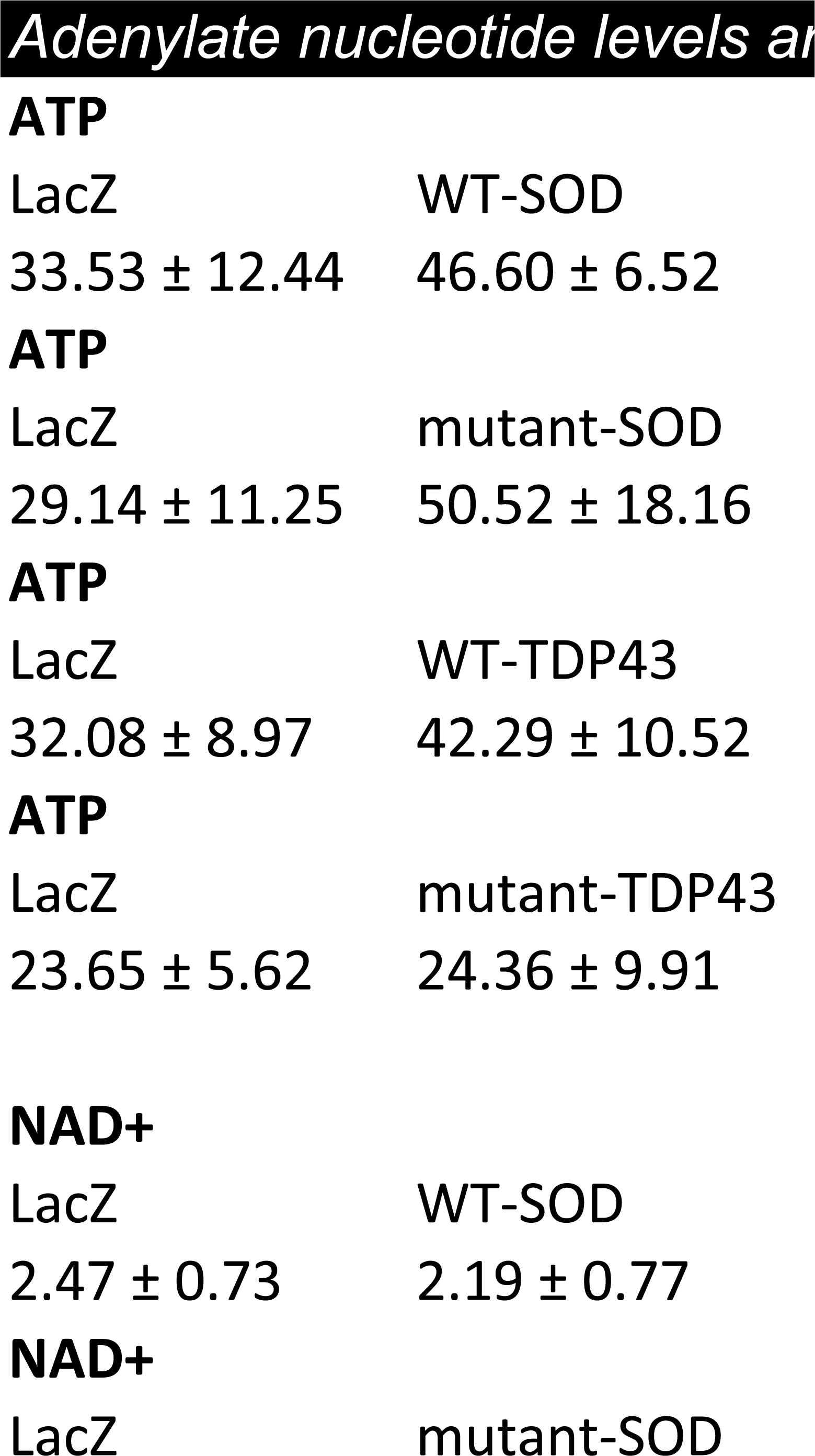
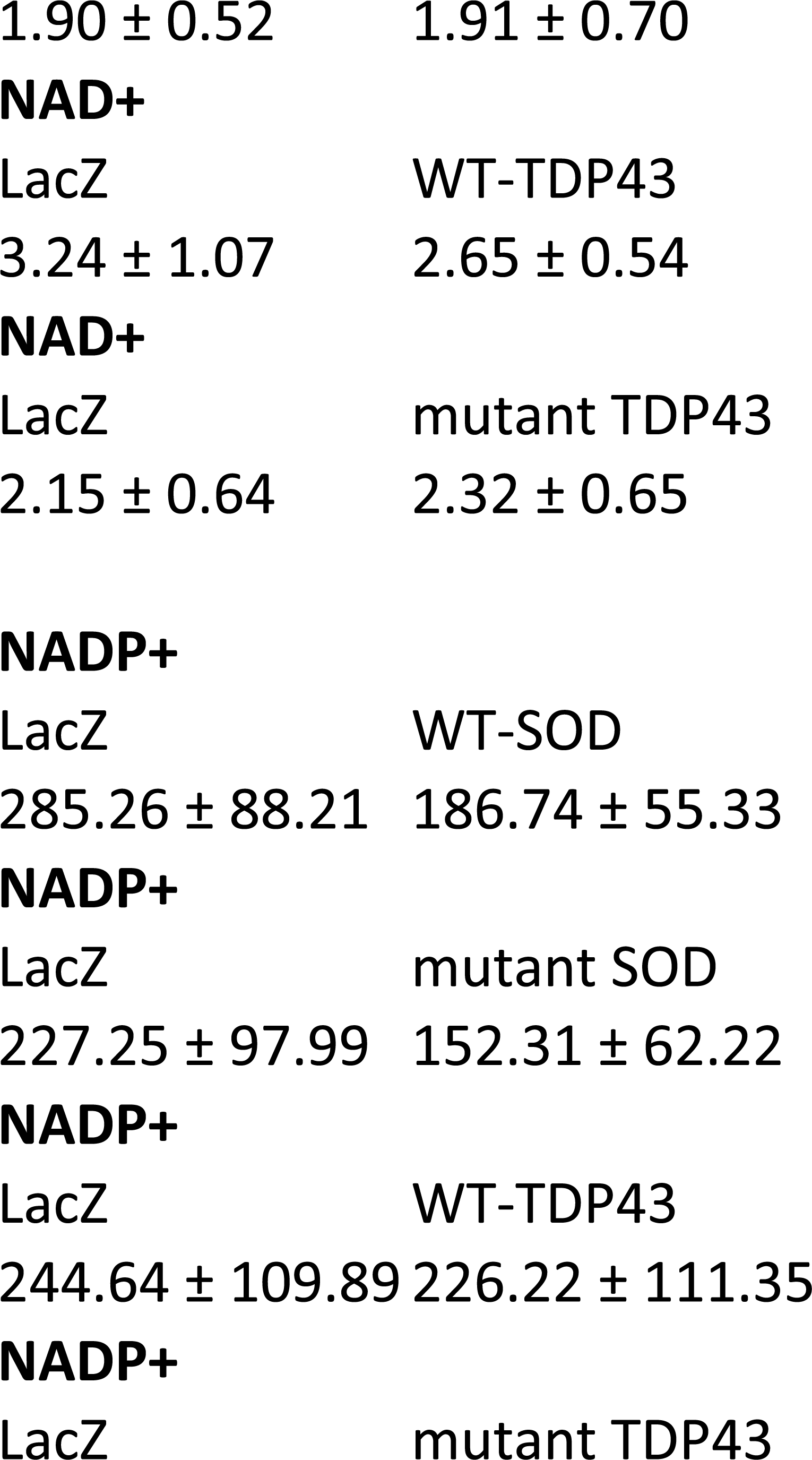
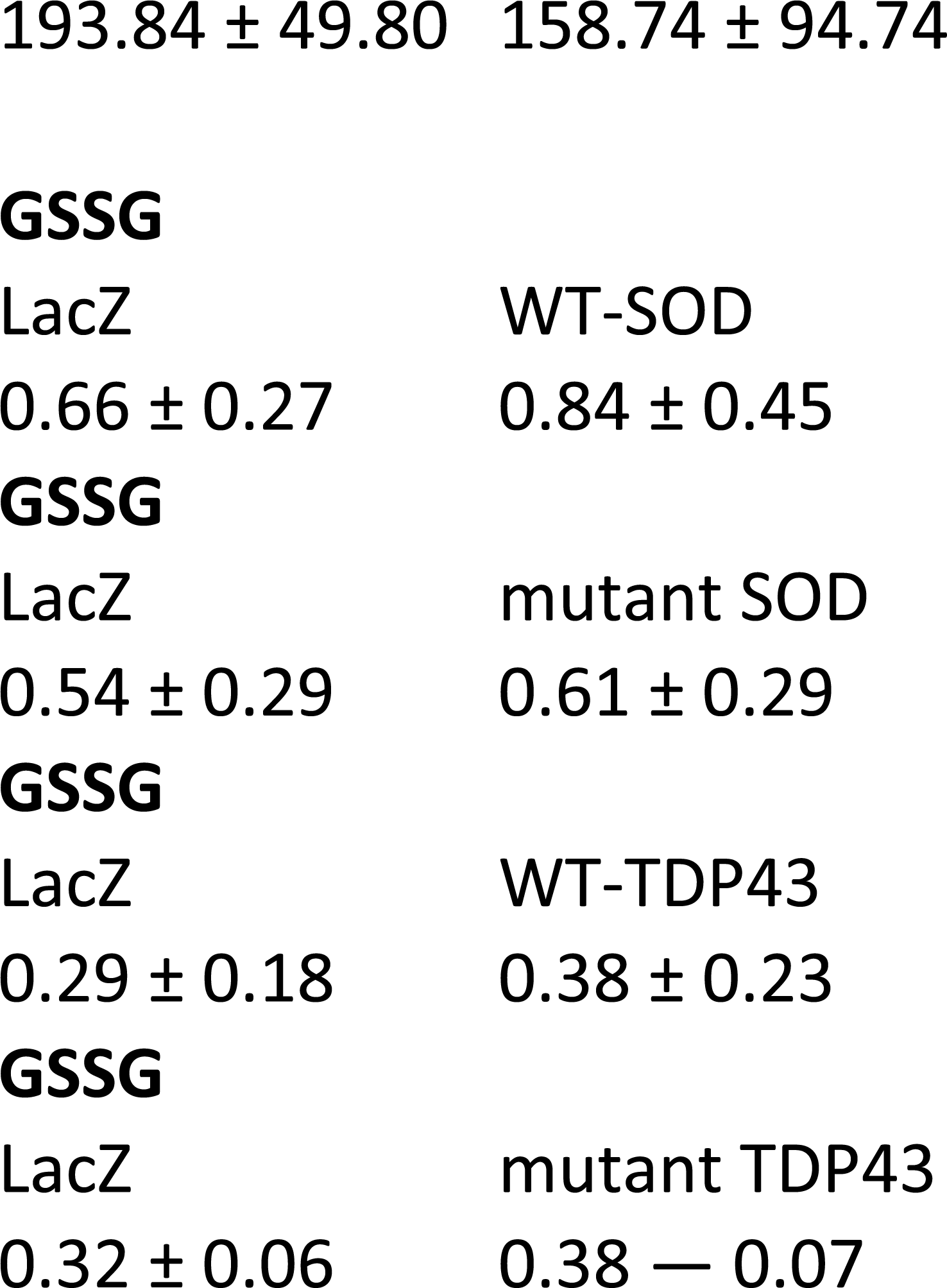
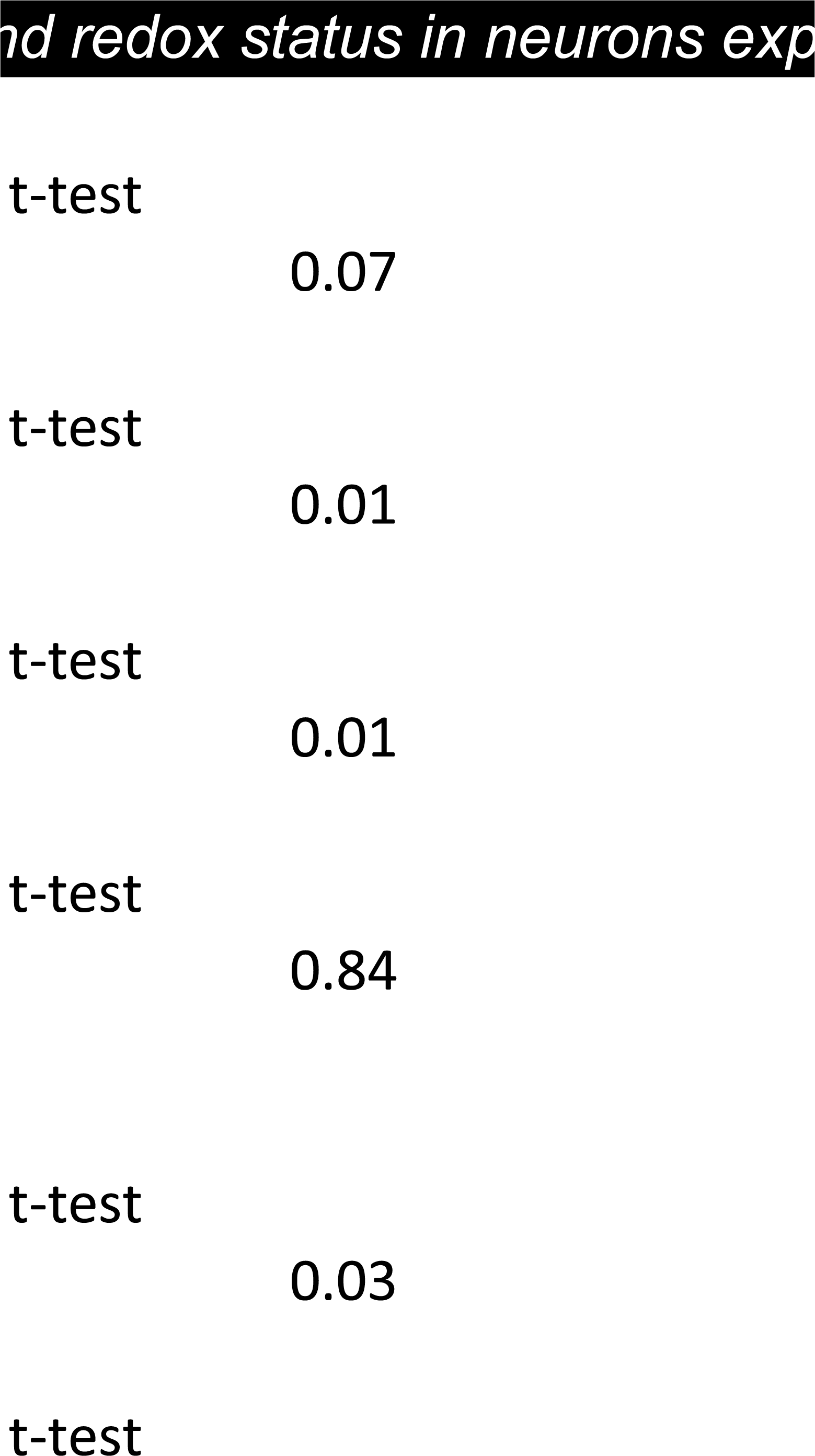
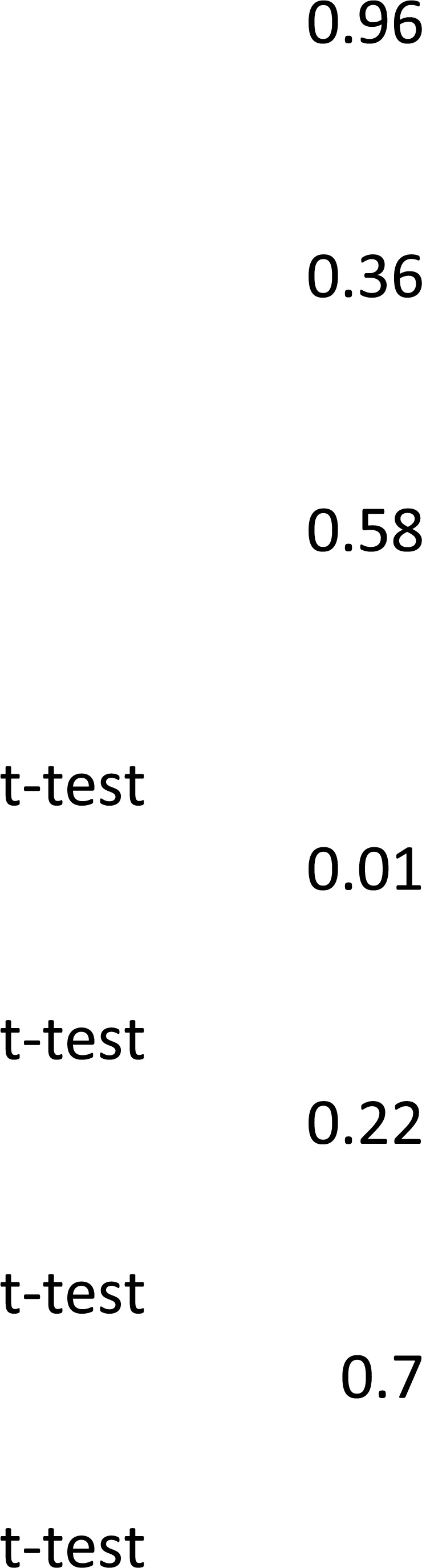
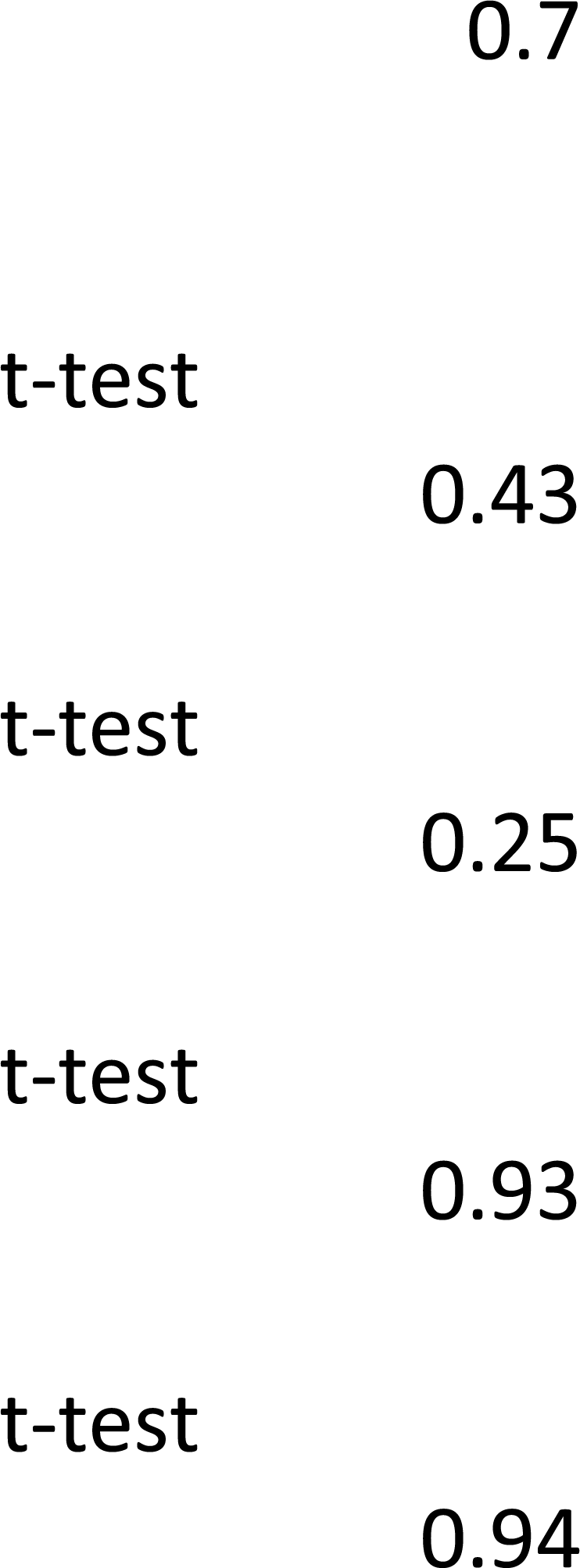
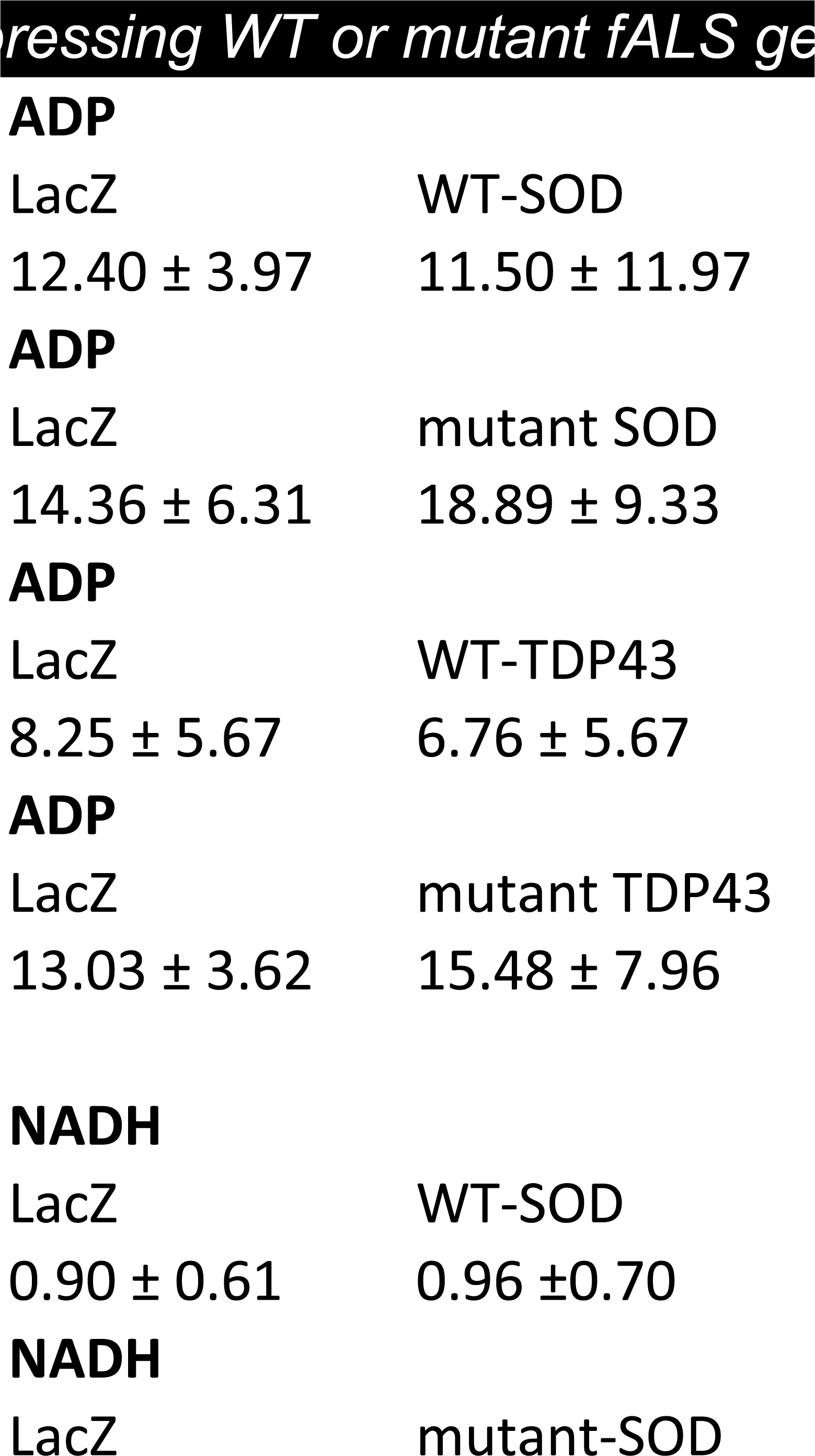
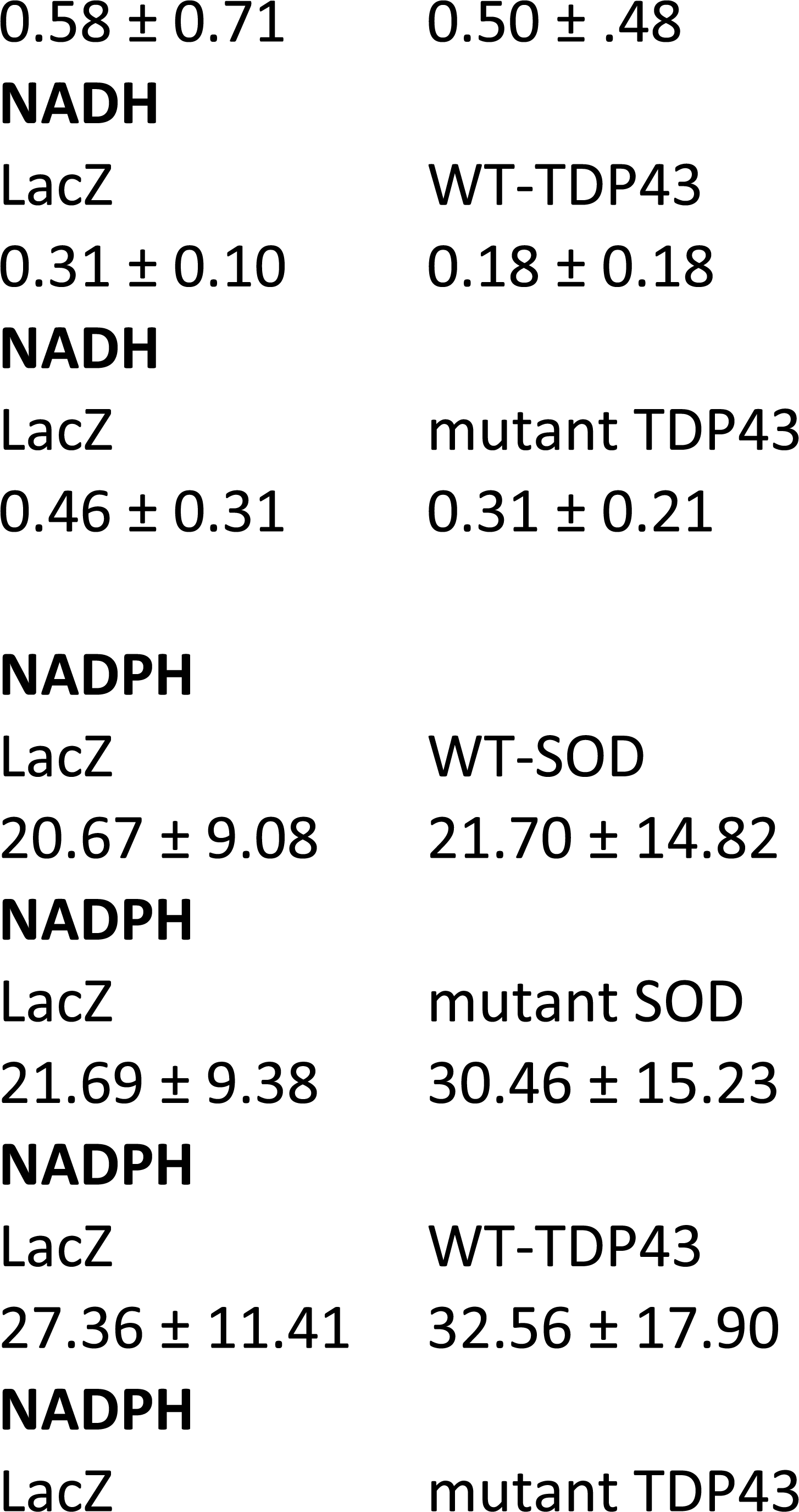
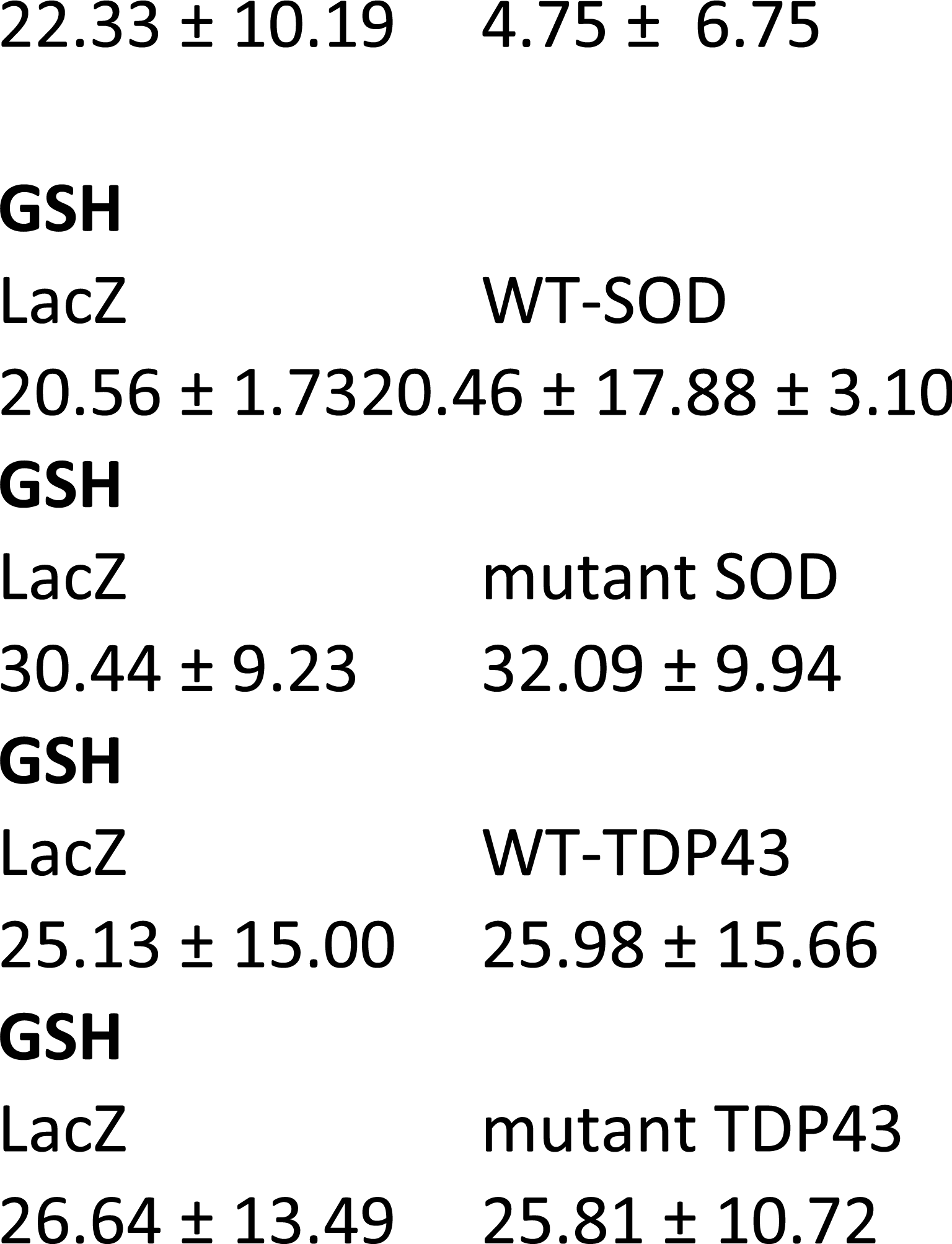
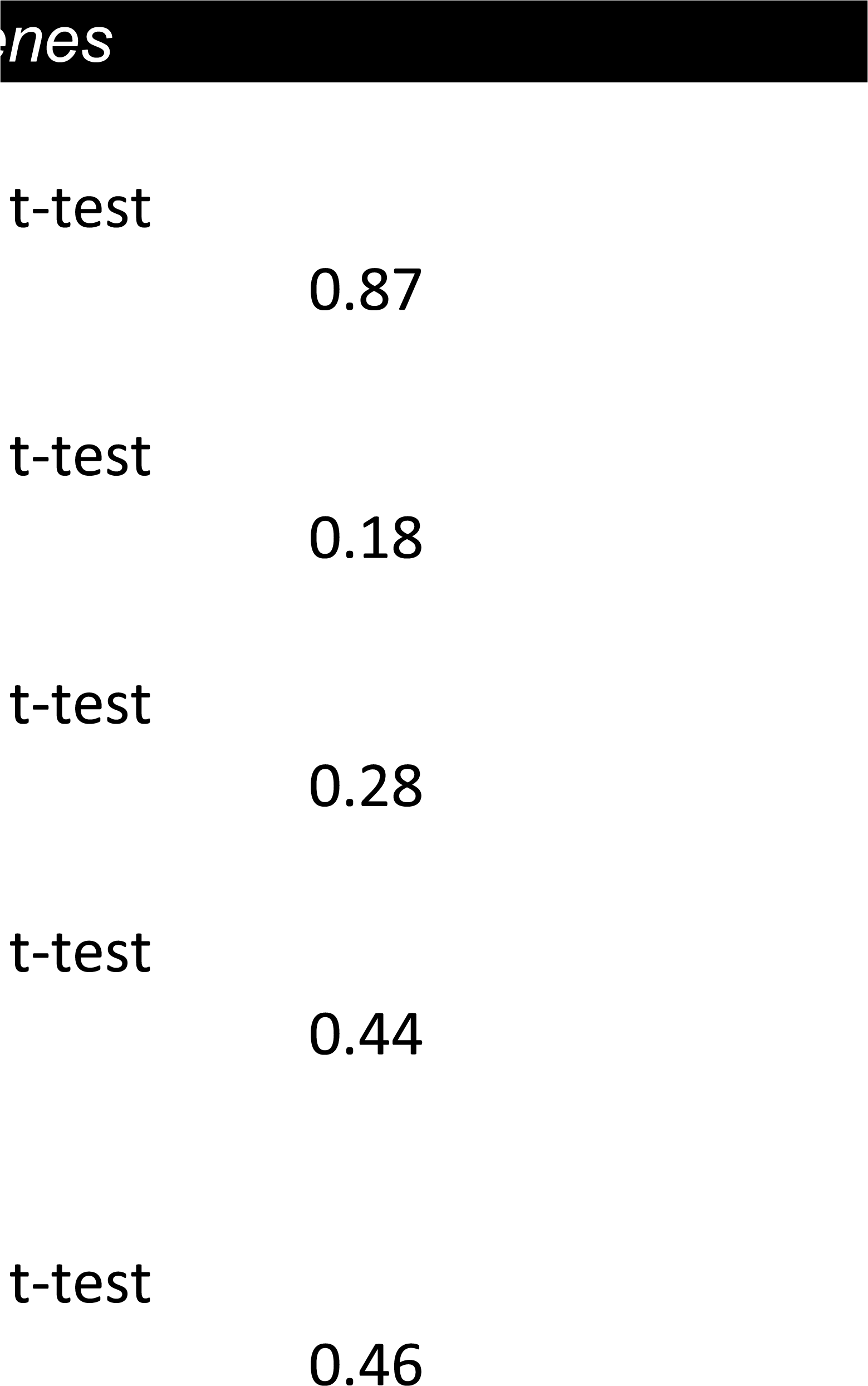
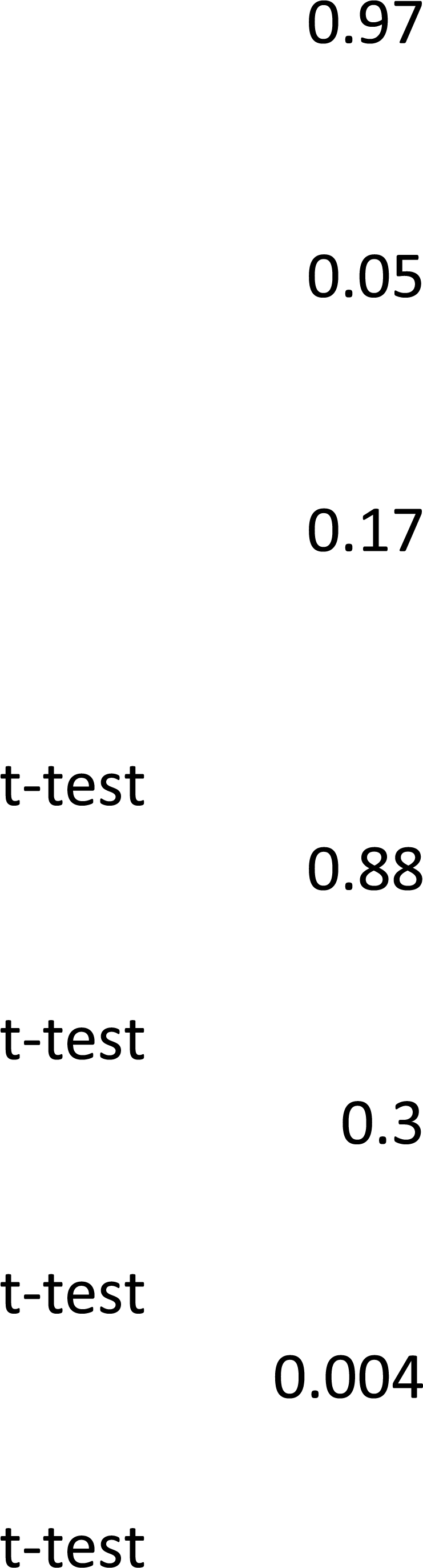
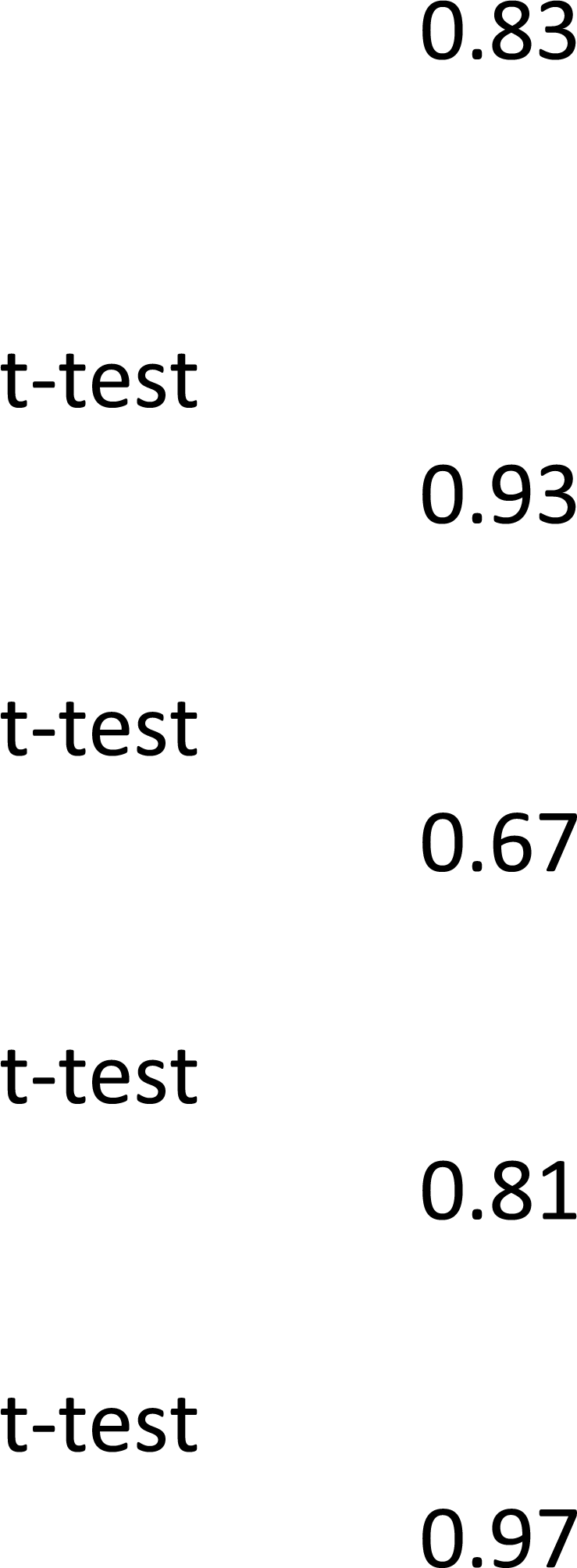
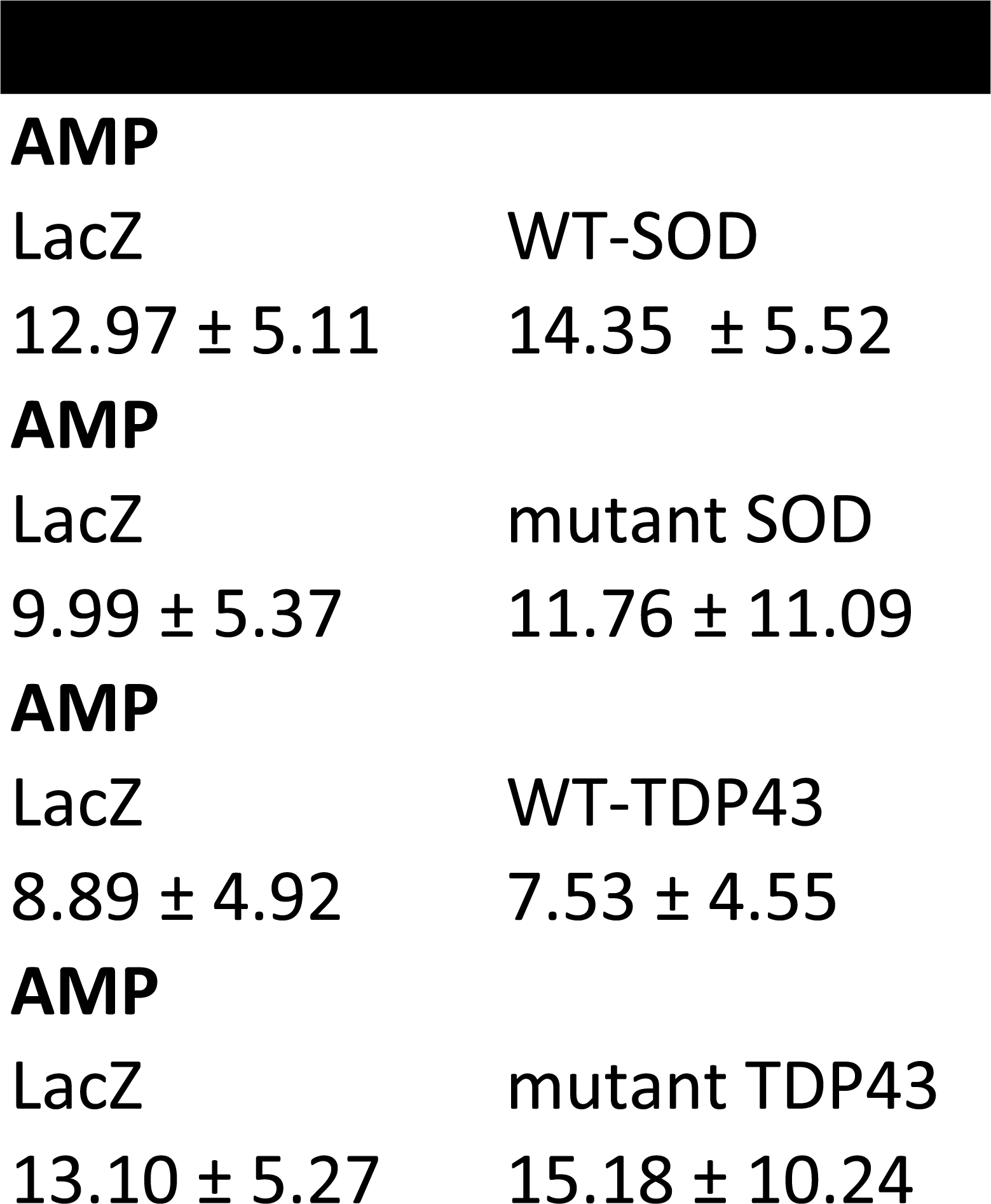
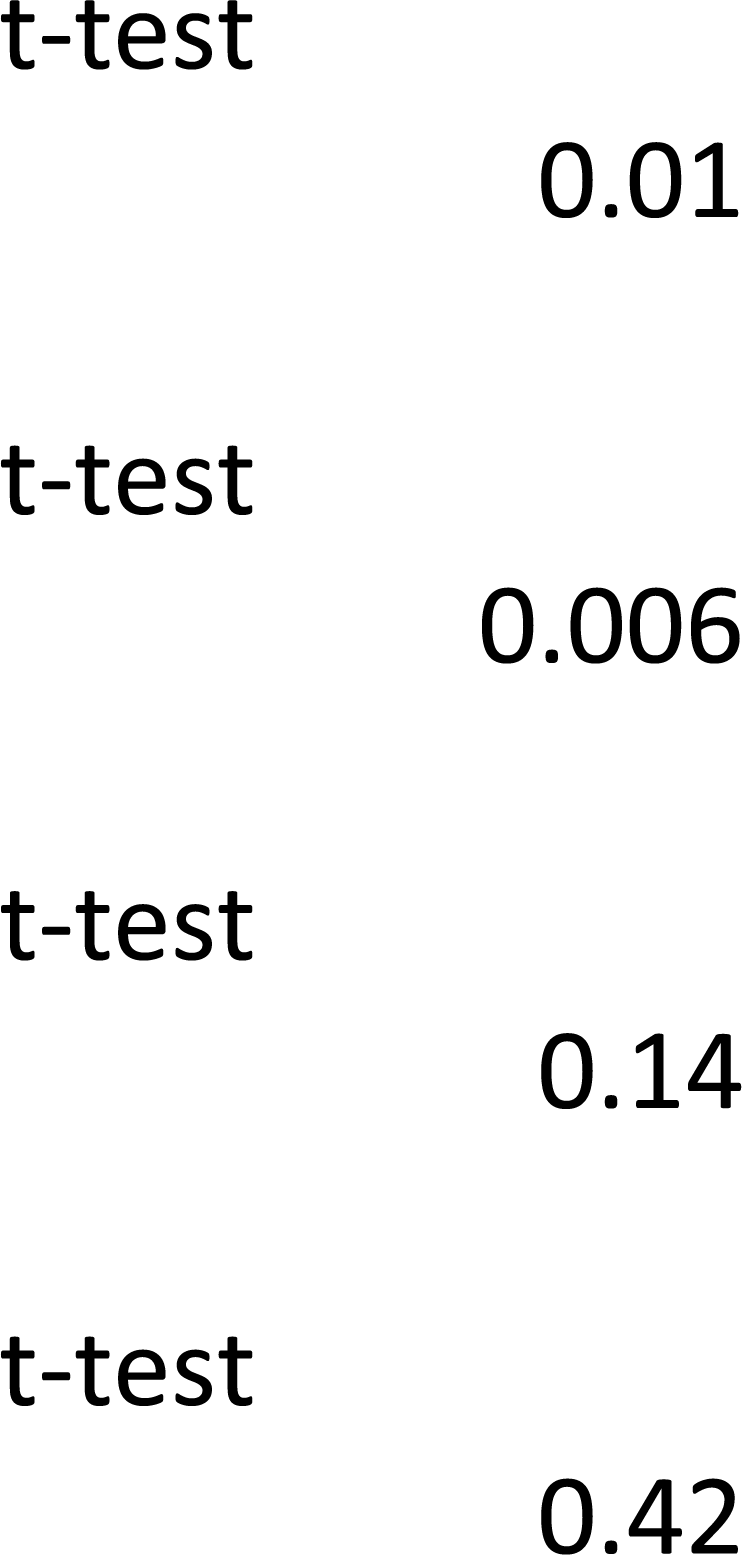
Adenylate nucleotide levels and redox status in neurons expressing Wild type or familial mutant ALS genes. Mean, standard deviation and Student’s t-test values from analysis of ATP/ADP/AMP, NAD+/NADH, NADP+/NADPH and GSSG/GSH are presented. Comparisons are from neurons expressing LacZ versus Wt-SOD1 versus Mt-SOD1 or LacZ versus Wt-TDP43 versus Mt-TDP43.

**Table 3.**
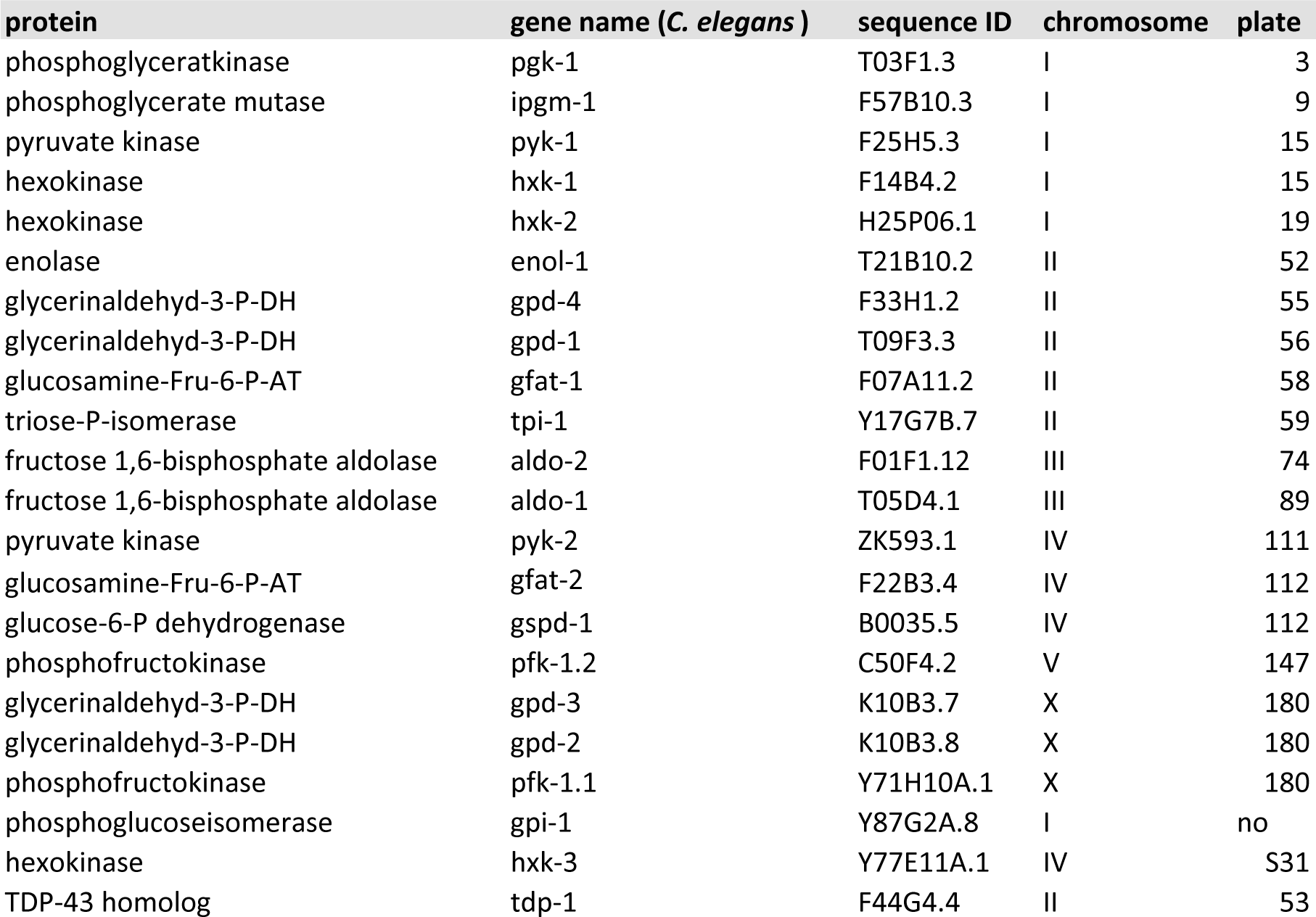
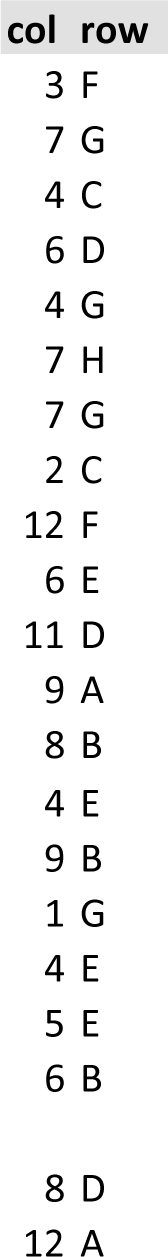
Yeast deletion strains used in these studies.

**Table 4.** Feeding RNAi bacteria studied in *C. elegans* model of fALS.

| protein | gene name (*C. elegans* ) | sequence ID | chromosome |
| --- | --- | --- | --- |
| glucosamine-Fru-6-P-AT | gfat-1 | F07A11.2 | II |
| glucosamine-Fru-6-P-AT | gfat-2 | F22B3.4 | IV |
| glucose-6-P dehydrogenase | gspd-1 | B0035.5 | IV |
| TDP-43 homolog | tdp-1 | F44G4.4 | II |

